# *Naegleria’s* mitotic spindles are built from unique tubulins and highlight core spindle features

**DOI:** 10.1101/2021.02.23.432318

**Authors:** Katrina B Velle, Monika Trupinić, Arian Ivec, Andrew Swafford, Emily Nolton, Luke Rice, Iva M. Tolić, Lillian K Fritz-Laylin, Patricia Wadsworth

## Abstract

*Naegleria gruberi* is a unicellular eukaryote whose evolutionary distance from animals and fungi has made it useful for developing hypotheses about the last common eukaryotic ancestor. *Naegleria* amoebae lack a cytoplasmic microtubule cytoskeleton and assemble microtubules only during mitosis, and thus provides a unique system to study the evolution and functional specificity of mitotic tubulins and the resulting spindle. Previous studies showed that *Naegleria* amoebae express a divergent α-tubulin during mitosis and we now show that *Naegleria* amoebae express a second mitotic α- and two mitotic β-tubulins. The mitotic tubulins are evolutionarily divergent relative to typical α- and β- tubulins, contain residues that suggest distinct microtubule properties, and may represent drug targets for the “brain-eating amoeba” *Naegleria fowleri*. Using quantitative light microscopy, we find that *Naegleria*’s mitotic spindle is a distinctive barrel-like structure built from a ring of microtubule bundles. Similar to those of other species, *Naegleria*’s spindle is twisted and its length increases during mitosis suggesting that these aspects of mitosis are ancestral features. Because bundle numbers change during metaphase, we hypothesize that the initial bundles represent kinetochore fibers, and secondary bundles function as bridging fibers.

## INTRODUCTION

Cells from across the eukaryotic tree use microtubules for a wide variety of functions during both interphase and mitosis. Interphase microtubules play essential roles establishing and maintaining cell shape, polarity, and intracellular trafficking. During cell division, a microtubule-based mitotic spindle self-assembles and mediates chromosome segregation. In most well-studied organisms, the spindle is composed of functionally distinct populations of microtubules, including: (1) kinetochore fiber microtubules that bind to kinetochores to connect each chromosome to a single spindle pole (Inoué and Salmon, 1995); (2) non-kinetochore microtubules that extend from the poles and overlap at the midzone, linking the two halves of the spindle (Mastronarde *et al.*, 1993; McIntosh, Molodtsov and Ataullakhanov, 2012); and (3) astral microtubules that extend from spindle poles toward the cell cortex. During anaphase, kinetochore microtubules shorten, while midzone microtubules elongate to drive chromosome segregation. A subset of midzone microtubules, called bridging fibers, contact kinetochore fibers in each half spindle (Kajtez *et al.*, 2016). Bridging fibers contribute to the balance of tension and compressive forces in the spindle (Kajtez *et al.*, 2016) and to chromosome motion in anaphase (Vukušić *et al.*, 2017; Vukušić, Buđa and Tolić, 2019). Spindle microtubules are organized by mitotic motor proteins that contribute to microtubule dynamic turnover, spindle pole organization, chromosome congression during prometaphase and poleward motion in anaphase. The influence of motor proteins in spindle structure is highlighted by the twist they introduce in spindles of human cells (Novak *et al.*, 2018).

Interphase and mitotic microtubule functions are emergent properties of microtubule-associated proteins as well as the subunit composition and post-translational modifications of the microtubule polymers themselves. Eukaryotic cells typically express multi-functional tubulins used for both interphase and mitotic functions (Raff, 1984). Human embryonic kidney cells, for example, express high levels of one α-tubulin and two 80% identical β-tubulins, which are used for both interphase and mitotic functions (Vemu *et al.*, 2017). Similarly, budding yeast express a single β-tubulin and two α-tubulins, which share 88% sequence identity and are used for both interphase and mitotic functions (Schatz et al., 1986). As an extreme example, the unicellular algae *Chlamydomonas* has a single α- and a single β-tubulin gene that are used for all microtubule functions (Johnson, 1998). Other eukaryotes, however, express unique tubulin isotypes that are required for specific microtubule functions, including meiotic spindle assembly in *Drosophila* oocytes (Matthews, Rees and Kaufman, 1993), axoneme formation in diverse systems (Hoyle and Raff, 1990), and touch receptor neurons in worms (Savage *et al.*, 1989). These specialized tubulins support the “multi-tubulin hypothesis” that posits that different tubulins can specify microtubules with distinct cellular functions (Wilson and Borisy, 1997).

The multi-tubulin hypothesis was inspired by studies of *Naegleria gruberi*—a single-celled eukaryote that diverged from the “yeast to human” lineage over a billion years ago (**Fig. 1A**)— with the unusual ability to differentiate from a crawling amoeba to a swimming flagellate (**Fig. 1B**) (Fulton and Simpson, 1976). The amoeba-to-flagellate differentiation is a stress response that involves the assembly of an entire microtubule cytoskeleton, including centrioles, flagella, and a complete cortical microtubule array. This process includes transcription and translation of flagellate-specific α- and β- tubulins along with their associated microtubule binding proteins (Fritz-Laylin, Assaf, *et al.*, 2010). The flagellate state is transient, and cells return to crawling amoebae within 2-300 minutes (Fulton, 1993), after which time the flagellate microtubules are disassembled and tubulin is degraded. This means that the *Naegleria* flagellate microtubules, and the α- and β- tubulins that comprise them, are specific for these non-mitotic microtubule functions.

**Figure 1.**
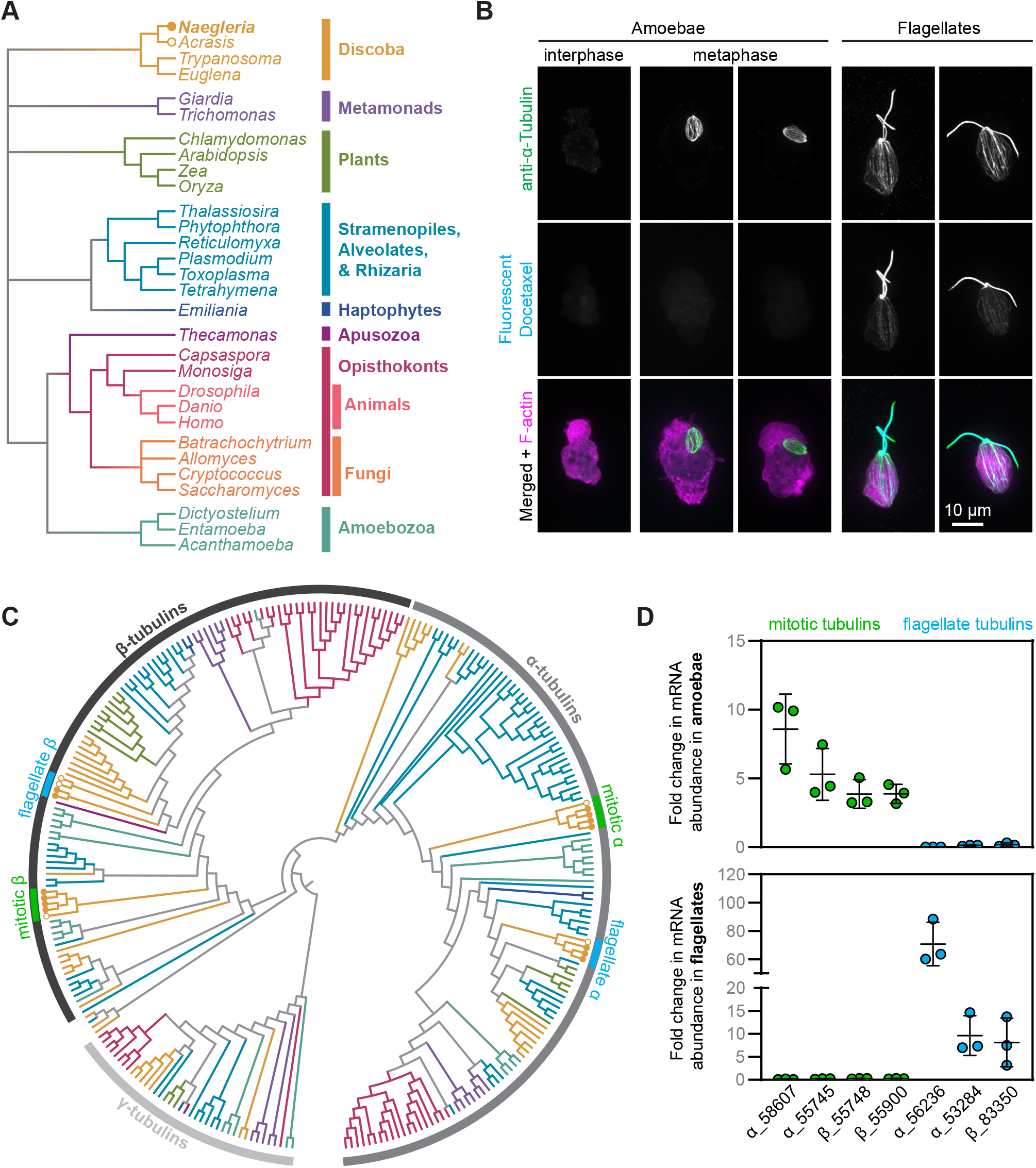
*Naegleria* has flagellate and mitotic microtubule arrays composed of distinct tubulins. (**A**) The evolutionary relationships between *Naegleria* and other eukaryotes are shown using a cladogram (branch lengths are meaningless) modified from Velle and Fritz-Laylin, 2019. (**B**) Amoebae from a growing population (left), or flagellates from a differentiated population (right), were fixed and stained with antibodies (anti-alpha tubulin clone DM1A, green) and Tubulin Tracker (Fluorescent Docetaxel, cyan) to detect microtubules, and Alexa Fluor 488 conjugated Phalloidin to label F-actin (magenta). Maximum intensity projections of cells are shown. (**C**) The evolutionary relationship of gamma, alpha, and beta tubulins from the species in panel A are shown using a cladogram (using the color scheme from A, see Fig. S1 for the full tree). The tree is rooted on gamma tubulins, and shows mitotic (green) and flagellate (blue) tubulins from *Naegleria* (closed circles) and *Acrasis* (open circles). (**D**) The fold changes in tubulin mRNA in amoebae compared to flagellates (top) or flagellates compared to amoebae (bottom) were calculated from data reported in Fritz-Laylin and Cande, 2010. Each point represents one experimental replicate, and lines denote the average +/− standard deviation (SD). Tubulins are labeled with JGI identification numbers.

In contrast to most eukaryotic cells, however, *Naegleria* amoebae have *no* observable interphase microtubules as visualized by immunofluorescence (**Fig. 1B**) (Walsh, 2007, 2012), or by electron microscopy (Fulton and Dingle, 1971). Moreover, interphase amoebae lack tubulin transcripts (Lee and Walsh, 1988; Chung *et al.*, 2002). Previous studies have shown that *Naegleria* has a divergent α-tubulin that is expressed specifically during mitosis (Chung *et al.*, 2002) and that is incorporated into its intra-nuclear mitotic spindle (Walsh, 2007, 2012). Because *Naegleria* uses this specific tubulin only for spindle assembly (Chung *et al.*, 2002), it provides a unique opportunity to examine a microtubule system that is specialized for mitosis. The *Naegleria* spindle also presents an interesting divergent morphology; instead of the typical rod- or fusiform-structures found in many eukaryotes, the *Naegleria* spindle is barrel-shaped and lacks both conventional kinetochores and obvious microtubule organizing centers (Fulton and Dingle, 1971; Akiyoshi and Gull, 2014; D’Archivio and Wickstead, 2017; Drinnenberg and Akiyoshi, 2017; van Hooff *et al.*, 2017).

Here we test whether—in the absence of the evolutionary constraints imposed by interphase microtubule functions—*Naegleria*’s mitotic microtubule system has diverged from canonical microtubule systems. We show that, in addition to the previously reported mitotic α-tubulin, *Naegleria* expresses a second mitotic α-tubulin along with two mitotic β-tubulins. In contrast to the *Naegleria* tubulins expressed during the flagellate stage that closely resemble tubulins from heavily-studied species, the protein sequences of the *Naegleria* mitotic tubulins have diverged significantly and have unique biochemical properties. We use quantitative microscopy to show that mitotic tubulins are used to build an unusual spindle composed of a ring of regularly-spaced microtubule bundles. As mitosis proceeds, additional microtubule bundles form in the equatorial region of the spindle and—as in other eukaryotes—the spindle elongates to facilitate chromosome segregation. The organization and dynamics of the *Naegleria* spindle highlight both core aspects of mitosis and as well variable features of cell division.

## RESULTS

### Naegleria expresses divergent α- and β-tubulins during mitosis

To determine the number and diversity of tubulins available to *Naegleria* amoebae and flagellates, we first searched for α- and β-tubulins in the *Naegleria gruberi* genome (Fritz-Laylin, Prochnik, *et al.*, 2010). As has been previously reported, we identified 13 α-tubulin and 9 β-tubulin genes, some of which appeared highly divergent, while others are closely related to those of other eukaryotes (Fritz-Laylin, Prochnik, *et al.*, 2010). To further explore the diversity of *Naegleria* tubulins, we reconstructed a maximum likelihood tree of α- and β-tubulins using γ-tubulins as an outgroup. Briefly, we collected and aligned 1,191 tubulins sequences from 200 different species (**Table S1, Datafile S1**), reconstructed a maximum likelihood tree (**Fig. S1, Datafile S2**), and pruned the resulting tree to more easily visualize the sequences of interest (**Fig. 1C, Datafile S3**). The tree recovers α-tubulins and β-tubulins as two, monophyletic clades with *Naegleria* mitotic and flagellar tubulin forming evolutionarily distinct clades within each tubulin family **(Fig. 1C)**.

The *Naegleria* α- and β-tubulin sub-clades most closely related to animal and fungal tubulins include those that are expressed during differentiation from the amoeba to the flagellate form (Lai *et al.*, 1979; Lee and Walsh, 1988; Fritz-Laylin and Cande, 2010). These tubulins represent the majority of axonemal and cytoplasmic tubulin protein in flagellates (Kowit and Fulton, 1974a, 1974b; Lai, Remillard and Fulton, 1988), and are not expressed in amoebae (Lai *et al.*, 1979; Lee and Walsh, 1988; Fritz-Laylin and Cande, 2010). Flagellate α-tubulins are 79-85% identical to human α-tubulin A1B (ENSP00000336799) and flagellate β-tubulins are 74-75% identical to human β-tubulin B1 (ENSP00000217133) **(Fig. S2C)**.

The second *Naegleria* tubulin sub-clades are more divergent. The second clade of α-tubulins contains two sequences from each *Naegleria gruberi* and *Naegleria fowleri,* and one from each of the related species *Acrasis kona, and Stachyamoeba lipophora*. The two *N. gruberi* α-tubulins are only 57-58% identical to human α-tubulin A1B **(Fig. S2C)**. Similarly, the second clade of *Naegleria* β-tubulins also includes *N. fowleri* and *A. kona* sequences, with *N. gruberi* sequences that are 57-58% identical to human β-tubulin B1.

Because the ortholog of the previously-reported mitotic α-tubulin (from the NB-1 strain) was among the divergent α-tubulins (from strain NEG-M) (Chung *et al.*, 2002; Fritz-Laylin, Prochnik, *et al.*, 2010), we predicted that the divergent *Naegleria* α- and β-tubulins are expressed during mitosis. Consistent with this prediction, we compared expression data of amoebae (a population that includes dividing cells) and flagellates and found the conserved tubulins expressed in flagellates and the divergent tubulins expressed in amoebae (**Fig. 1D**). We confirmed this finding by comparing expression levels of the putative-mitotic tubulins in mitotically synchronized cells to control cell populations and found at least two-fold enrichment of the divergent tubulin transcripts (**Fig. S2A-B**). Together these data indicate that *Naegleria gruberi* amoebae expresses divergent α- and β-tubulins during cell division.

### *Naegleria* mitotic tubulins have diverged in ways that suggest distinct biochemical properties

Visual inspection of *Naegleria* mitotic and flagellate tubulin sequences suggested that the mitotic tubulins may have altered microtubule dynamics and/or binding sites for microtubule associated proteins. To more systematically assess this possibility, we quantified the divergence of mitotic and flagellate α- and β-tubulins as a function of amino acid position. Briefly, after building master multiple sequence alignments for α- and β-tubulins containing mitotic and flagellate tubulin sequences from *N. gruberi, N. fowleri,* and *A. kona* along with reference sequences from more commonly studied organisms (see Methods), we made separate ‘mitotic’ and ‘flagellate’ subalignments for each species by only retaining the mitotic or flagellate tubulin from that species (in addition to the reference sequences). We used these subalignments to measure the difference in conservation at each position, and we summarized the results with a positional ‘divergence score’ (**Fig. 2A**) in which negative values correspond to greater divergence at a given amino acid position (see Methods). Mitotic α-tubulins have more positions with elevated divergence compared to β-tubulin in all three species (compare **Fig. 2A** top and bottom), although the absolute number of divergent positions differs by organism (35 positions in α-tubulin vs 23 in β-tubulin for *N. gruberi*; 24 vs 22 for *N. fowleri*; 32 vs 27 for *A. kona*).

**Figure 2.**
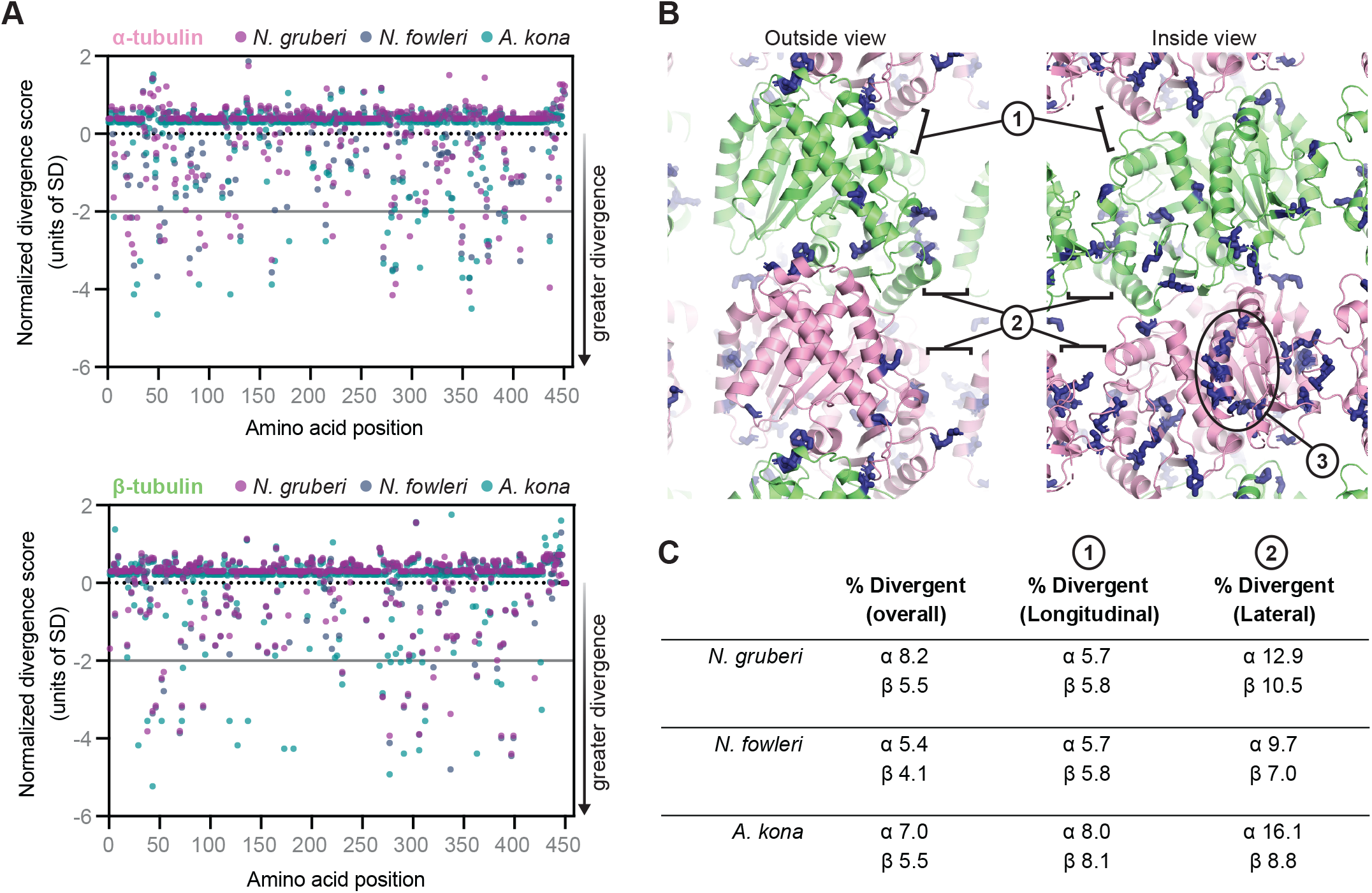
Comparative analysis of evolutionary divergence for mitotic and flagellate tubulins. (**A**) Plots of the normalized divergence score (see Methods) as a function of amino acid position for α-tubulin (top) and β-tubulin (bottom). Lower scores indicate positions where mitotic tubulins show increased divergence relative to flagellate tubulins. The analysis was performed on three species: *N. gruberi* (lavender), *N. fowleri* (navy), and *A. kona* (teal). The horizontal gray line indicates the two standard deviation cutoff we used to identify especially divergent sites. (**B**) Structural context of the sites with increased divergence in the mitotic tubulins. Side-chain positions for the *N. gruberi* amino acids identified in (A) are represented as sticks (blue) on a model of αβ-tubulin in the microtubule lattice (α-tubulin: pink, β-tubulin: lime). ‘Outside’ and ‘Inside’ views of the lattice are shown, and longitudinal (labeled 1) and lateral (labeled 2) microtubule lattice contacts are indicated, as is the luminal (internal) surface of α-tubulin (labeled 3). (**C**) Table summarizing the proportion of positions with elevated divergence near microtubule lattice interfaces. For all three species, there are more divergent positions in α-tubulin compared to β-tubulin, and the divergence seems to be particularly enriched at the lateral interfaces. See Fig. S4 for details.

Although the positions of elevated variability are distributed throughout the tubulin fold for both α- and β-tubulin, they appear to be enriched near microtubule polymerization interfaces and surfaces displayed on the inside of the microtubule (**Fig. 2B, Fig. S3**). To quantify this impression, we tested for enrichment at longitudinal or lateral polymerization interfaces by determining whether the fraction of divergent positions near a given interface was greater than the fraction of divergent positions across the entire sequence (see Methods). This analysis reveals that divergent positions are more enriched at lateral lattice contacts (2-3-fold increase depending on the species) than at longitudinal lattice contacts (1.1-1.9-fold, depending on the species; **Fig. 2C**). This enrichment of divergence at lattice interfaces reinforces the idea that microtubules formed from mitotic tubulins will have altered polymerization dynamics and/or distinct structural features.

Because fluorescent docetaxel—a microtubule labeling reagent derived from the microtubule-stabilizing drug taxol—appears to bind *Naegleria* flagellate tubulin but not mitotic tubulin (**Fig. 1B**), we next examined if taxol-binding residues were conserved in either of these sequences. We focused our analysis on β-tubulin sequences because that is where taxol binds, and we selected taxol-binding residues based on prior analyses (Gupta *et al.*, 2003) and the structure of taxol-bound microtubules (Alushin *et al.*, 2014). Important taxol-binding amino acids are conserved in flagellate but not in mitotic β-tubulin sequences (**Fig. S4A**). Thus, consistent with our observation that fluorescent docetaxel only labels flagellate microtubules, flagellate tubulins appear to have an intact taxol binding site, whereas the mitotic tubulins appear to have lost the ability to bind taxol.

Finally, we noted interesting sequence differences in disordered regions of the *Naegleria* tubulins. For example, the major site of α-tubulin acetylation, K40, is conserved in the flagellate tubulins, but has diverged in the mitotic tubulins (**Fig. S4B**). We also characterized the length and predicted net charges of the C-terminal tubulin tails (**Fig. S4C**). The tubulin tails of both mitotic and flagellate α-tubulins have lengths and net changes similar to those observed in more commonly studied tubulins. In contrast, the mitotic β-tubulin tails are slightly less charged than their flagellate counterparts (**Fig. S4C**). Finally, the C-terminal EY sequence in α-tubulin that is recognized by regulatory factors that contain a CAP-GLY domain is notably absent from both flagellate and mitotic tubulin sequences, suggesting differences in their regulation (**Fig. S4C**). Together, these observations reinforce the notion that microtubules assembled from mitotic αβ-tubulins are likely to have different polymerization dynamics and/or binding partners compared to microtubules assembled from flagellate αβ-tubulins.

### The *Naegleria* spindle is a hollow barrel of microtubule bundles that elongates as mitosis proceeds

To explore whether the sequence divergence of *Naegleria*’s mitotic tubulins translates into a divergent organization of mitotic microtubules, we fixed *Naegleria* amoebae undergoing closed mitosis. We stained mitotic microtubules with anti-tubulin antibodies and DNA with DAPI, and visualized the cells using spinning disk confocal microscopy (**Fig. 3**). Consistent with previous results (Fulton and Dingle, 1971; Chung *et al.*, 2002; Fritz-Laylin *et al.*, 2011; Walsh, 2012), we find that the *Naegleria* spindle is composed of microtubule bundles and lacks obvious microtubule organizing centers (**Fig. 3**). The microtubule bundles appear to form around a ball of DNA; we refer to this stage as prophase (**Fig. 3A**). This cage-like array of microtubule bundles reorganizes into a barrel-shaped spindle with DNA aligned in a broad, hollow band at the midplane; we refer to this stage as metaphase (**Fig. 3A**). Although in some cases the spindle has a tapered morphology (**Fig. 3A**, left metaphase cell) the majority of spindles are characterized by broad, flat poles (**Fig. 3A**, middle and right metaphase cells). We also observed spindles in which the DNA is segregated to the ends of the elongated spindle, which we classified as anaphase/telophase. Compared with other stages of mitosis, few spindles were detected during the early stages of chromosome segregation, suggesting that this stage occupies a small fraction of the total duration of mitosis. In contrast, cells with elongated spindles and segregated DNA were relatively common, suggesting that the late anaphase spindle is stable for some time. By quantifying spindle length and width, we infer that spindle length increases while width decreases as mitosis progresses from prophase to anaphase/telophase (**Fig. 3B**).

**Figure 3.**
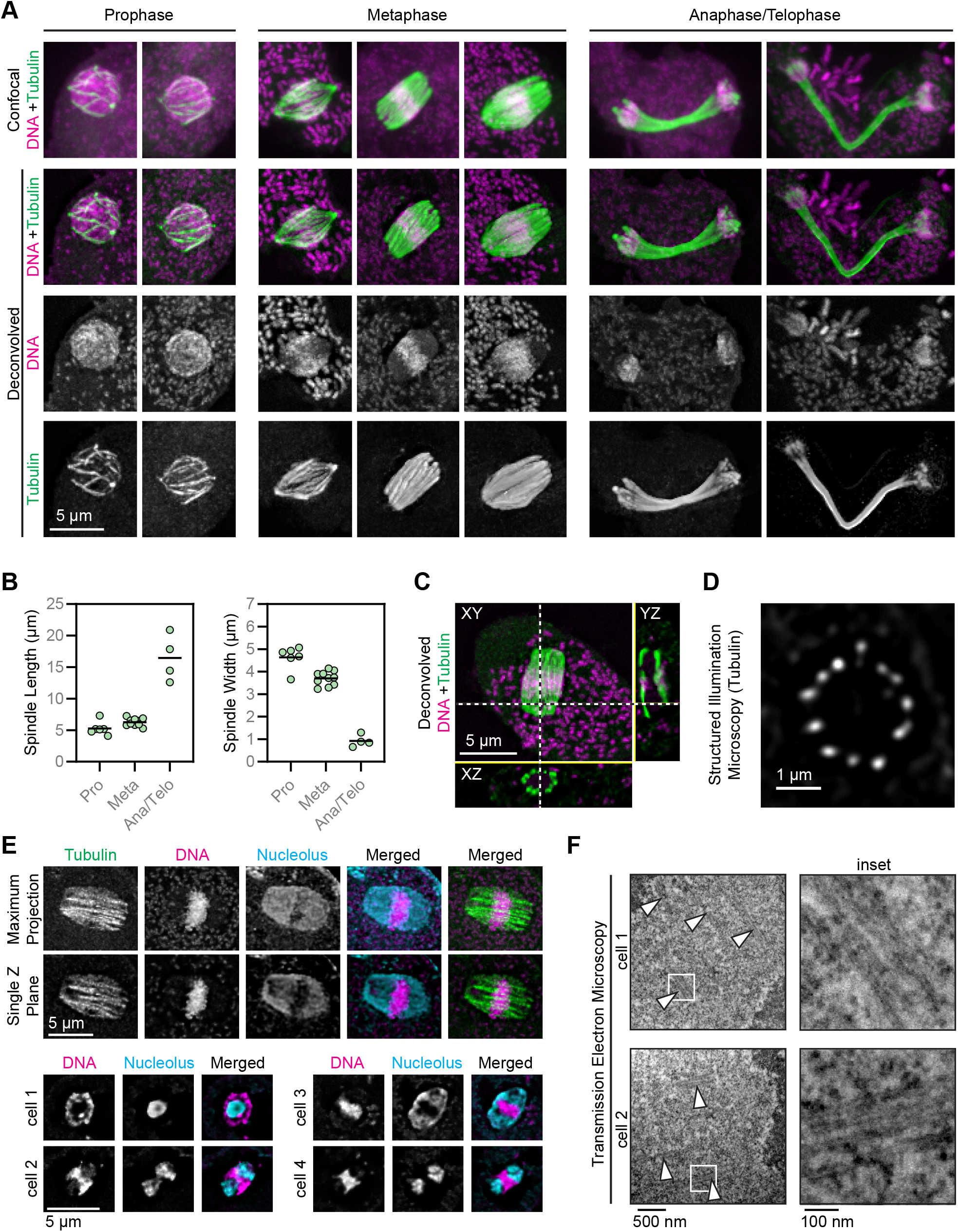
*Naegleria*’s spindle is a barrel shape composed of bundles of microtubules that elongate as mitosis proceeds. (**A**) Asynchronously growing *Naegleria* amoebae were fixed and stained with anti-alpha tubulin clone DM1A (green) to detect microtubules, and DAPI to label DNA (magenta). Mitotic spindles were imaged using confocal microscopy (top row), and images were deconvolved using Autoquant software (bottom rows). Cells were classified as prophase, metaphase, or anaphase/telophase. (**B**) Quantification of maximum spindle length (left) and the spindle width at half the length (right). Each point represents one mitotic spindle, and lines indicate the averages (prophase, n=6; metaphase, n=10; anaphase/telophase, n=4). Spindles imaged and deconvolved as in (A). (**C**) Orthogonal views of a metaphase spindle (imaged and deconvolved as in A) lying in the plane of the coverslip; XZ and YZ views generated in Fiji. (**D**) Structured illumination microscopy of a spindle lying perpendicular to the coverslip. (**E**) Confocal microscopy and deconvolution of nucleoli in mitotic *Naegleria*. Cells were fixed and stained to detect tubulin (YOL 1/34 antibody, green, top panels only), DNA (DAPI, magenta), and nucleolar protein (DE6 antibody, cyan). One maximum intensity projection is shown (top cell), while remaining images are single Z planes. (**F**) Transmission electron microscopy of microtubule bundles in *Naegleria*; arrowheads indicate microtubule bundles and boxed regions (left) are shown as enlarged insets (right).

Because mitotic cells were relatively rare in asynchronous populations, we also examined mitotically synchronized cells (Fulton and Guerrini, 1969) (**Fig. S5**), and found no qualitative or quantitative differences in spindle microtubule organization between the synchronized and asynchronous cells (**Fig. S5C**). This supports previous reports that synchronization does not alter spindle morphology in *Naegleria* amoebae (Fulton and Guerrini, 1969). We therefore used cells from both synchronized and asynchronized populations for the following analyses.

To determine the organization of microtubule bundles in the *Naegleria* spindle, we visualized axial and transverse slices of spindles oriented both parallel and perpendicular to the coverslip (**Fig. 3C-D**). These analyses confirmed that the microtubules in the *Naegleria* metaphase spindle are organized in a ring, similar to the staves of a barrel (**Fig. 3C-D**). Previous studies have suggested that this barrel is assembled around the nucleolus, which remains intact during mitosis (*Naegleria*’s ribosomal RNA genes are encoded on a plasmid that does not condense during prophase (Fritz-Laylin *et al.*, 2011; Walsh, 2012)). To confirm the retention of the nucleolus during mitosis, we co-stained cells with anti-nucleolar and/or anti-tubulin antibodies, as well asDAPI to visualize DNA (**Fig. 3E)**. Consistent with previous work, we find that the nucleolus remains throughout mitosis, at times encompassing much of the spindle volume (Walsh, 2012). The nucleolus divides before chromosome segregation, resulting in one nucleolus at each end of the spindle with the chromosomes nestled between them in a thin disk (**Fig. 3E**).

Comparing the dimensions and intensity of the microtubule arrays in flagellates to those in mitotic cells suggests that the spindle is composed of bundles rather than individual microtubules (**Fig. 1B**). Supporting this idea, we observed a single anaphase cell in which a microtubule bundle appears to have splayed apart, revealing at least five fluorescent elements which may represent individual microtubules (**Fig. S5E**). To estimate the number of microtubules per bundle, we fixed *Naegleria* amoeba for thin section transmission electron microscopy. Longitudinal sections through mitotic cells reveal that bundles are composed of multiple, closely-associated individual microtubules (**Fig. 3F**). We observed three to six microtubules in a single longitudinal section consistent with previous estimates in the related *N. fowleri (González-Robles et al., 2009)*. In summary, our data show that the *Naegleria* spindle is composed of a ring of microtubule bundles that elongates during chromosome segregation.

### *Naegleria* spindles have two sets of microtubule bundles

Although most spindles were oriented parallel to the coverslip surface, some spindles were oriented perpendicular to the coverslip, providing improved resolution of the microtubule bundles (**Fig. 4A**). These end-on views revealed variation in the number of microtubule bundles (**Fig. 4A-C**). Some spindles have a single ring of approximately 12 evenly-spaced bundles with 0.79 μm center-to-center spacing (range: 0.42-1.90; SD: 0.28; n: 31 measurements from 3 spindles). These “primary bundles” extend the entire length of the spindle (**Fig. 4A,** left, **Fig. 4B**, top). Other spindles, however, have additional bundles adjacent to the main ring (**Fig. 4A**, middle and right, **Fig. 4B** bottom). Importantly, the number of bundles in this second class of spindles varied along the spindle axis, with additional “secondary bundles” restricted to the spindle midplane with the primary bundles extending out to the spindle ends.

**Figure 4.**
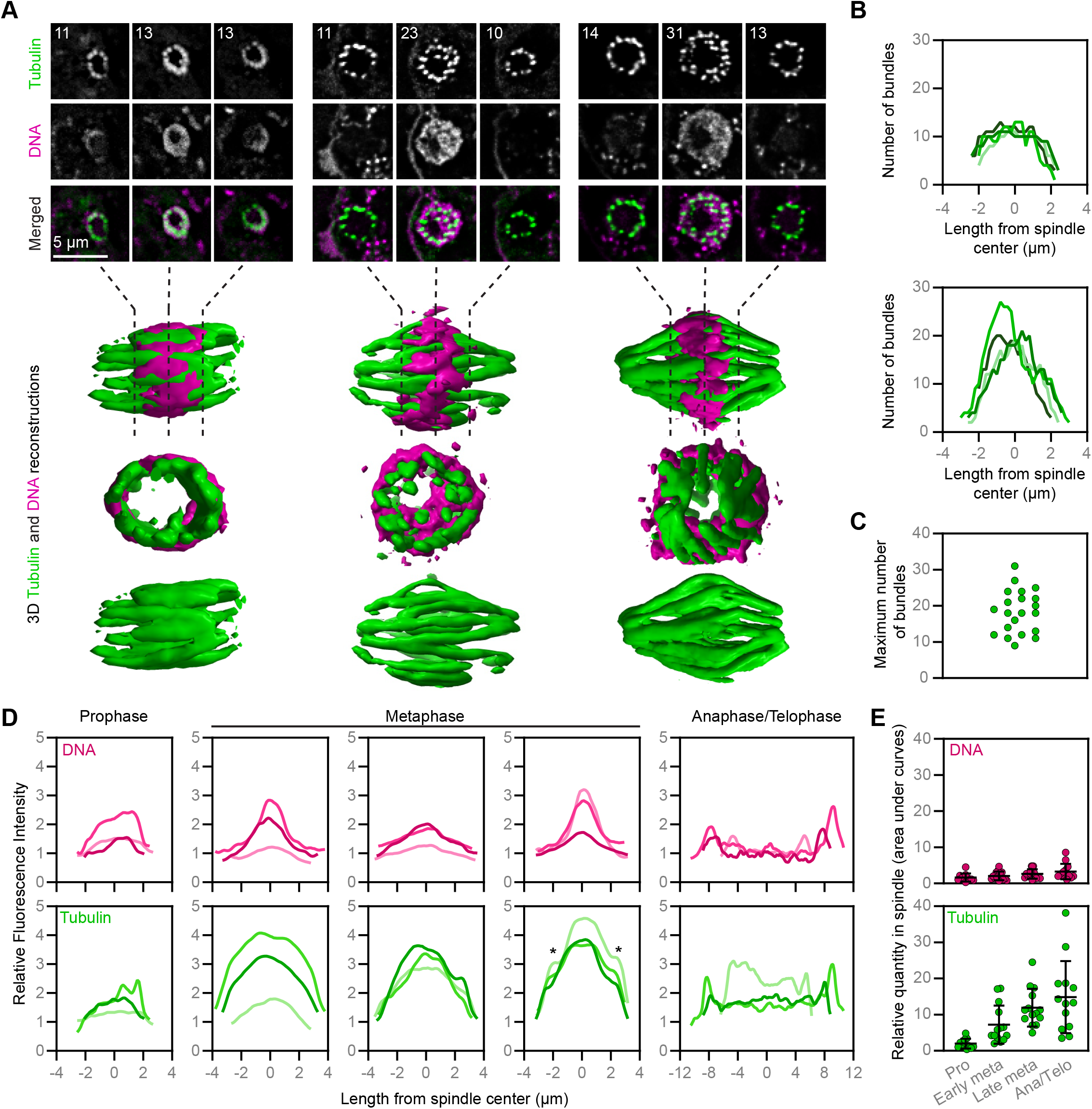
The number of microtubule bundles changes as mitosis proceeds. (**A**) Cells were fixed and stained with antibodies (anti-alpha tubulin clone DM1A, green) to detect microtubules, and DAPI to label DNA (magenta). Cells with spindles perpendicular to the coverslip were imaged using confocal microscopy and deconvolved using Autoquant software (top panels), and 3D reconstructions were rendered using ChimeraX software (bottom panels, not to scale). Individual Z planes are shown for slices approximately 25, 50, and 75% through the spindle for three representative cells. Numbers (upper left) indicate the number of distinct microtubule bundles in that position of the spindle. (**B**) The number of microtubule bundles throughout the spindle length in metaphase spindles, imaged as in (A). Some spindles (top) had a fairly consistent number of microtubule bundles throughout the spindle (n=4), while other spindles (bottom) had a peak in the number of bundles towards the midpoint (n=4). (**C**) The maximum number of microtubule bundles from confocal images of metaphase cells. (**D**) Line scans show the relative DNA and tubulin fluorescence intensity from sum intensity projections of spindles lying in the plane of the coverslip, imaged as in (A). Metaphase spindles were grouped based on the shapes of tubulin curves (no shoulders, left; unclear shoulders, center; two clear shoulders denoted by asterisks, right); three individual examples are shown in each panel (also see Fig. S6). (**E**) Quantification of DNA (top) or tubulin (bottom) from line scans obtained as in (D). Metaphase was categorized as early or late based on the presence (late) or absence (early) of shoulders (stages where no clear classification could be assigned were excluded). Each point represents the area under the curve for one spindle line scan, and lines indicate the mean +/− SD.

If the secondary bundles were formed from new microtubule polymerization, we would expect the mid-region of metaphase spindles to have a greater amount of tubulin than the poles. We therefore quantified tubulin and DNA fluorescence intensity along horizontally-oriented spindles at each stage of mitosis (**Fig. 4D, Fig. S6**). The total amount of tubulin within the spindle increases as mitosis proceeds, consistent with microtubule assembly (**Fig 4E**). Metaphase spindles show variable tubulin distributions (**Fig. 4D**), with a subset having a clear peak of intensity toward the spindle midzone with “shoulders” on either side (**Fig. 4D,** rightmost metaphase). This pattern is reminiscent of the larger number of bundles that we quantified at the centers of vertically oriented spindles (**Fig. 4B**), and is consistent with secondary bundle formation involving additional microtubule assembly. Although this subset of metaphase spindles had clear “shoulders” in their tubulin distributions, other distributions were less clear-cut (**Fig. 4D,** center metaphase panel). The variability in the tubulin distribution across metaphase spindles raises the possibility that secondary bundles may form asynchronously within a spindle, consistent with cross sections of vertically-oriented spindles that show only a few secondary bundles (**Fig. 4A**, middle cell). By quantifying the maximum number of bundles per vertically-oriented spindle (**Fig. 4C**), we found that the maximum bundle number varies from ~10 to 25, with many cells showing intermediate values. This continuous distribution is consistent with asynchronous secondary bundle assembly rather than the two distinct populations we would expect for a synchronous event.

Finally, we examined the tubulin distribution in anaphase and telophase cells to determine the fate of the secondary bundles that form during metaphase. Although the tubulin intensity in these spindles was relatively uniform across the spindle midzone, we observed distinct peaks at each end of the spindle, indicating a higher density of microtubules (**Fig. 4D,** anaphase), consistent with both primary and secondary bundles remaining associated with chromosomes throughout mitosis. Together, these data suggest that secondary microtubule bundles assemble asynchronously during metaphase by new microtubule assembly and may persist through late mitosis. Based on these and other data, we hypothesize that primary bundles serve as kinetochore fibers and secondary bundles as bridging fibers (see Discussion).

### The *Naegleria* spindle twists from pole-to-pole in a right-handed fashion

The 3D reconstructions of vertically-oriented spindles revealed that the microtubule bundles curved and appeared to twist from one end of the spindle to the other (**Fig. 4A**, **Fig. 5A**, **Movie S1, Movie S2**). To quantify this, we traced individual bundles of metaphase spindles (**Fig. 5A**) and measured their curvature and twist by fitting a plane to the points representing the bundle and a circle that lies in this plane to the same points. We then estimated bundle curvature as one over the radius of the fit circle, and the twist as the angle between the plane and the z-axis divided by the mean distance of these points from the z-axis (**Fig. 5B**).

**Figure 5.**
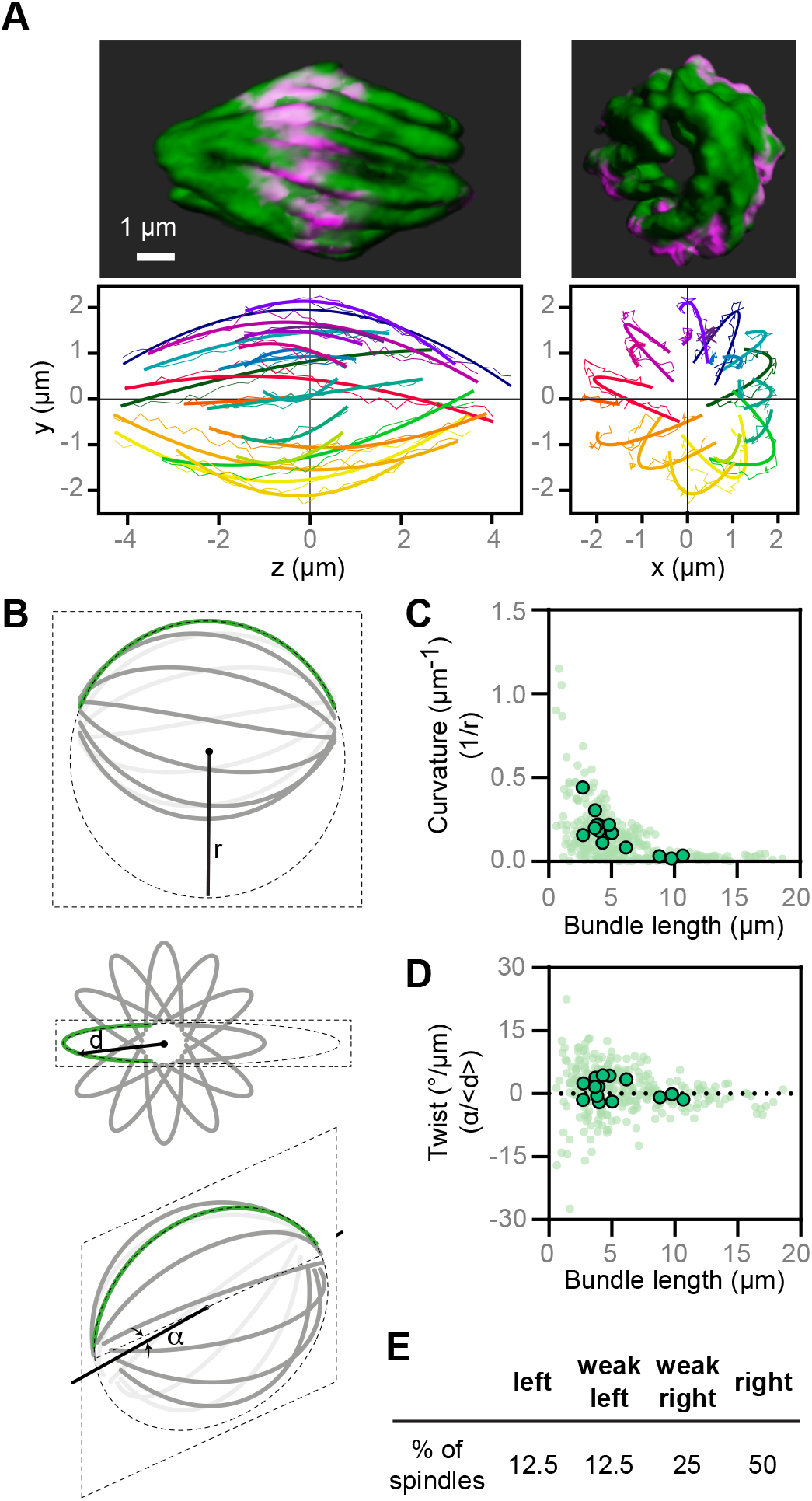
*Naegleria* mitotic spindles are twisted. (**A**) A 3D reconstructed spindle (the same spindle shown in Fig. 4A, right) is shown from side and end-on view viewpoints. Microtubules are shown in green, and DNA is in magenta. Microtubule bundles were quantified from the side view (left graph) and end-on view (right graph). Each bundle is represented by a different color, thin lines mark the manually traced points along the bundle, and thick lines show circular arcs of the fitted circles. (**B**) A simplified scheme of a spindle is shown from the side (top), end-on (middle), and from an arbitrary angle (bottom). A microtubule bundle (green line) is fitted by a circle (dashed ellipse) of radius (r). The angle (α) between the central spindle axis (solid line) and the plane in which the fitted circle lies (dashed parallelogram) is denoted. The distance (d) of the bundle from the central spindle axis is denoted. (**C**) The curvature of microtubule bundles is shown as a function of bundle length (measured along its pole to pole axis). Each small dot represents a single bundle within a spindle, while each larger dot represents the average for a spindle. (**D**) The twist of microtubule bundles is shown as a function of bundle length. Each small dot represents a single bundle within a spindle, while each larger dot represents the average for a spindle. (**E**) The percentage of spindles with right, weak right, left, or weak left handedness are shown (see Fig S7 for a breakdown of this analysis).

The resulting data show that microtubule bundles in the *Naegleria* spindle are curved (0.146 ± 0.009/μm, **Fig. 5C**) and twisted (0.873 ± 0.316 degrees/μm; positive values denote right-handed and negative values left-handed twist **Fig. 5D**), with shorter bundles having more curve and twist than longer bundles (**Fig. 5C-D**). On average, the bundles were twisted in a right-handed direction, making the spindle a chiral structure with right-handed asymmetry. This result was corroborated by visual assessment of the handedness of the spindle twist. Here, if the bundles rotate counterclockwise when moving along the spindle axis in the direction towards the observer, the twist is right-handed, and vice versa. We found a mixture of left- and right-handed twist, with the majority of spindles showing a strong right-handed twist (**Fig. 5E**). Analyzing early metaphase (defined as cells with <20 bundles) and late metaphase (defined as cells with >20 bundles) cells separately suggests that bundles increase in length and decrease in curvature during metaphase (**Fig. S7A and S7D**). Right-handed twist was dominant for vertically- and horizontally-oriented spindles, and for cells in early and late metaphase (**Fig. S7B and G**), suggesting that the handedness of spindle chirality does not depend on mitotic stage or spindle orientation during imaging. Together, these data indicate that the microtubule bundles are physically linked and under rotational forces.

Other than HeLa cells, *Naegleria* are the only cell type whose spindle twist has been measured. The microtubule bundles of *Naegleria*’s spindle are less curved than those of human HeLa cell spindles, as the radius of curvature is larger for *Naegleria*, 6.9 ± 0.4 μm, than for the outermost bundles in HeLa cells, 5.1 ± 0.3 μm (Manenica *et al.*, 2020). Moreover, the radius of curvature normalized to the spindle half-length, which is equal to 1 for bundles shaped as a semicircle, is 1.26 ± 0.05 for *Naegleria* and 0.90 ± 0.05 for HeLa cells (Manenica *et al.*, 2020), also indicating a smaller curvature of Naegleria spindles. In line with the smaller curvature, the absolute value of the average spindle twist in *Naegleria* is smaller than in HeLa cells, 0.9 ± 0.3 degrees/μm in *Naegleria* vs. 2 degrees/μm in HeLa (Novak *et al.*, 2018). Yet, twist of *Naegleria* spindles is more eye-catching than in HeLa cells, due to the smaller number of microtubule bundles, which are well-defined and have a uniform shape, in contrast to the less ordered distribution and shapes of bundles in HeLa cells.

## DISCUSSION

*Naegleria* amoebae represent a remarkable system with which to study microtubule biology because they do not have interphase microtubules. *Naegleria* is not the only species without interphase microtubules; the cytoplasm of interphase *Entamoeba histolytica* amoebae also has no observable microtubules (Meza, Talamás-Rohana and Vargas, 2006). In contrast to *Entamoeba,* however, *Naegleria* can differentiate into a secondary cell type, the flagellate. Here we show that *Naegleria* express unique tubulins in mitotic amoebae that are distinct from the tubulins expressed in flagellate cells. While flagellate tubulins—used to assemble both flagellar and cytoplasmic microtubules (Fulton and Kowit, 1975; Fulton and Simpson, 1976; Fulton, 1983; Lai, Remillard and Fulton, 1988; Fritz-Laylin and Cande, 2010)—are highly similar to tubulins of other eukaryotic species, the mitotic tubulins have diverged in sequence, including at key residues likely to alter microtubule structure or dynamics. Because the sequence similarity between *Naegleria* and *Acrasis* flagellate tubulin isoforms is much higher than their mitotic tubulins (Fig. S2-C), we infer that the cytoplasmic functions of tubulins may require more stringent sequence conservation than mitotic functions.

*Naegleria* mitotic microtubules assemble into a hollow, barrel-shaped mitotic spindle comprising distinct bundles, each made of multiple microtubules. Based on these observations and additional literature discussed below, we propose the following model for *Naegleria* spindle elongation and chromosome segregation (**Fig. 6**): (1) Mitosis begins with the assembly of “primary” microtubule bundles. Each primary bundle is associated with a chromosome and functions as a pair of kinetochore fibers; (2) During metaphase, “secondary” microtubule bundles form near the spindle midplane that function as bridging fibers, connecting kinetochore fibers associated with sister chromatids (Simunić and Tolić, 2016; Vukušić *et al.*, 2017); (3) chromosome-to-pole motion occurs as primary bundles depolymerize while secondary bundles elongate to form the spindle midzone and further separate the chromosomes. Under this model, the higher microtubule density toward the poles during late anaphase results from fluorescence of both kinetochore and bridging fiber microtubules. While this model is consistent with our quantitative measurements, other scenarios are also possible. For example, secondary bundles could associate with chromosomes, functioning like kinetochore fibers, and primary bundles could form the midzone, or each individual bundle could be composed of bridging and kinetochore fiber microtubules, that ultimately sort into the anaphase spindle.

**Figure 6.**
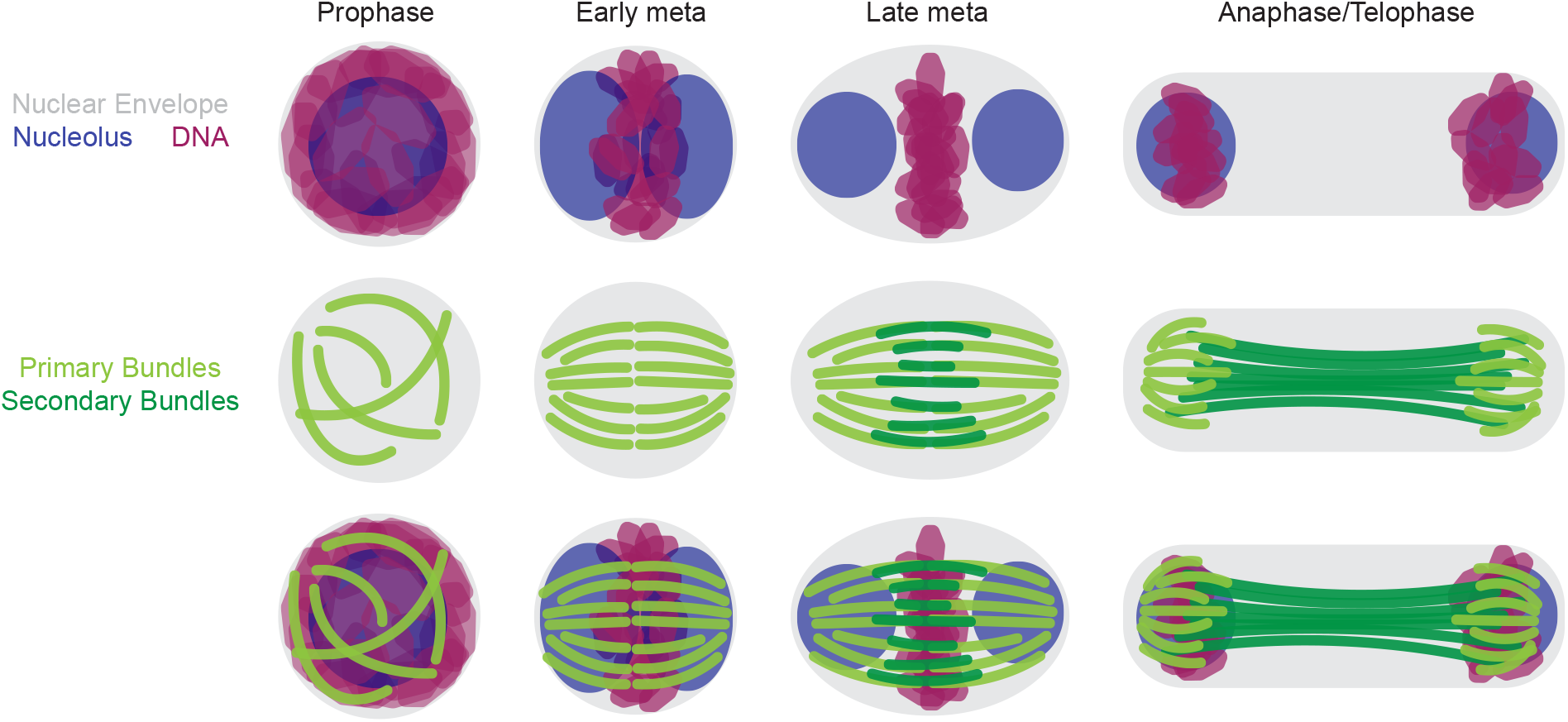
Model for mitosis in *Naegleria*. During prophase in *Naegleria*, bundles of microtubules form around a hollow sphere of DNA (magenta) which surrounds the single, round nucleolus (blue). In early metaphase, the DNA condenses into a disk, the nucleolus begins to divide and the microtubule bundles (light green) organize into a hollow, twisted barrel shape. In late metaphase, the DNA is further condensed, and the nucleolus resolves into two distinct spheres. A secondary set of microtubules forms in the equatorial region (dark green) adjacent to the primary bundles. During anaphase/telophase, the DNA is segregated to the two ends of the spindle and the spindle elon-gates. See text for details.

The possibility that primary bundles function as kinetochore fibers is consistent with our previous estimate of ~12 chromosomes in *Naegleria* (Fritz-Laylin, Prochnik, *et al.*, 2010), a value that is similar to the average number of primary bundles we observe in early metaphase spindles (**Fig. 4**). We also observe “kinks” in the center of some spindles suggesting that each primary bundle may be composed of two kinetochore fibers. Although conventional, trilaminar kinetochores have not been detected using electron microscopy (**Fig. 3F**) (Fulton and Dingle, 1971), homologs of a subset of kinetochore proteins identified in other organisms are present in the *Naegleria* genome (Akiyoshi and Gull, 2014; van Hooff *et al.*, 2017), hinting at the presence of yet-to-be-detected kinetochores. Whether or not *Naegleria* has conventional kinetochores, spindle assembly and chromosome movement is well established to occur in the absence of kinetochores (Heald *et al.*, 1996; Brunet *et al.*, 1999). For example, in both mouse and *C. elegans* meiotic and human mitotic spindles, lateral interactions between microtubules and chromosomes drive chromosome congression, although chromosome-to-pole motion does require kinetochore-microtubule interactions (Kapoor *et al.*, 2006; Mullen, Davis-Roca and Wignall, 2019; Danlasky *et al.*, 2020).

Our working model posits that both anaphase A, chromosome-to-pole motion, and anaphase B spindle elongation, contribute to chromosome segregation in *Naegleria*. The presence of short microtubule bundles between the chromosomes and poles in anaphase spindles is consistent with microtubule depolymerization during anaphase A, although the location and regulation of microtubule assembly and disassembly in these cells is not yet known. Anaphase/telophase spindles in *Naegleria* are longer than metaphase spindles, consistent with anaphase B spindle elongation. In meiotic spindles, anaphase B is driven, at least in part, as polymerizing midzone microtubules interact with chromosomes (Dumont, Oegema and Desai, 2010; Danlasky *et al.*, 2020). In mammalian cells, links between elongating midzone bridging microtubules and kinetochore fibers contribute to anaphase (Vukušić *et al.*, 2017; Vukušić, Buđa and Tolić, 2019). Although the mechanism of spindle elongation in *Naegleria* is not yet established, the appearance of secondary bundles in the chromosome region is reminiscent of bridging fibers in other cell types (Simunić and Tolić, 2016). This similarity suggests that interactions between primary and secondary microtubule bundles may contribute to chromosome segregation (Vukušić, Buđa and Tolić, 2019).

These bundles differentiate the *Naegleria* spindle from those of other species that typically contain both individual and bundled microtubules. Despite this difference, microtubule bundles in *Naegleria* and human cells both show twist, suggesting that this may be a conserved feature of eukaryotic spindles. In contrast to the left-handed chirality previously measured in human spindles (Novak *et al.*, 2018), the majority of *Naegleria* spindles are right-handed. Because *Naegleria* is only the second species whose spindle chirality has been measured, it is difficult to know whether its chirality is unusual. Regardless, the requirement of the motor activity of kinesin Eg5 in the twisting of human spindles suggests that *Naegleria* spindle twist may also depend on the activity of microtubule motors that generate torque within the bundles (Tolić, Novak and Pavin, 2019).

*Naegleria*’s evolutionary position makes it uniquely suited for identifying features of mitotic spindles that may be deeply conserved, including their bi-polarity, elongation, and twist. *Naegleria*’s position also highlights features that may be lineage-specific due to their absence in this distant species. For example, some features of animal cell spindles are missing from *Naegleria,* including obvious microtubule organizing centers as well as astral microtubules which contribute to spindle position and to cytokinesis in other cells. Whether these differences are related to the divergence of the *Naegleria* mitotic tubulins awaits further investigation.

The unique properties of these mitotic tubulins may also have practical value. Although the model species *Naegleria gruberi* is innocuous, the related *Naegleria fowleri* is the infamous “brain-eating amoeba” that causes a devastating and usually lethal brain infection (Siddiqui *et al.*, 2016). Because the divergent residues we have identified in the *Naegleria* mitotic tubulins are conserved in both *Naegleria* species but not in human tubulins (**Fig. 2, Fig. S4**), these residues represent potential targets for specific therapeutics that could disrupt *Naegleria* cell division to halt *in vivo* growth.

## MATERIALS AND METHODS

### Phylogenetic tree estimation

To establish a more inclusive comparison of *Naegleria* α-, and β-tubulins to those of other eukaryotes, 1,191 tubulins from 200 different species were analyzed (Table S1), adding sequences from *Naegleria gruberi (Fritz-Laylin, Prochnik, et al., 2010)*, *Naegleria fowleri* (Herman *et al.*, 2020), and *Acrasis kona* (personal communication, Sandra Baldauf, Uppsala University) to those identified as α, β, and γ tubulins using the PhyloToL pipeline (Cerón-Romero *et al.*, 2019). Prior to alignment, sequences from the same species that were 100% identical were removed, leaving only one copy before re-merging the datasets. Sequences were aligned using the PASTA iterative alignment algorithm with the MUSCLE algorithm as the aligner and merger (Mirarab *et al.*, 2015). IQ-Tree v1.16.2 was used for model selection, which indicated LG4M+R10 as the best model for reconstruction (Kalyaanamoorthy *et al.*, 2017; Minh *et al.*, 2020). Due to the size of the tree, LG4M was used balance the accuracy of tree solving and the constraints of modern processing power. A maximum likelihood tree was reconstructed using IQ-Tree with 10,000 ultrafast bootstraps (Hoang *et al.*, 2018). 1,000 bootstraps of the approximate likelihood ratio test (Guindon *et al.*, 2010) as well as the aBayes test (Anisimova *et al.*, 2011) were then used to further test node support. The ITOL web server was used for tree visualization (Letunic and Bork, 2019).

### Characterization of *Naegleria* mitotic tubulin sequences

To quantify the divergence of mitotic and flagellate α- and β-tubulins from *N. gruberi*, *N. fowleri*, and *A. kona* as a function of amino acid position, we compared them to a common reference consisting of sequences of α- or β-tubulin sequences from commonly studied model organisms (*Homo sapiens, Sus scrofa, Bos taurus, Drosophila melanogaster, Mus musculus, Saccharomyces cerevisiae, Schizosaccharomyces pombe*, and *Chlamydomonas reinhardtii*). Multiple sequence alignments were first prepared for α- and β-tubulin using ClustalOmega (Madeira *et al.*, 2019). These ‘master’ alignments contained the reference sequences as well as mitotic and flagellate sequences from the three species of interest. Separate “flagellate” and “mitotic” subalignments were then prepared for each species by only retaining flagellate or mitotic sequences from a given species, in addition to the common reference sequences. We quantified sequence conservation/divergence as a function of amino acid position in these subalignments using the AL2CO server (Pei and Grishin, 2001), using normalized sum of pairs scoring (BLOSUM62 weighting) and otherwise default settings. The resulting conservation scores are normalized so that completely conserved positions return the same score regardless of the identity of the conserved amino acid; lower scores (including negative scores) correspond to less conservation. To assess differences in conservation between mitotic and flagellate sequences, the flagellate score was subtracted from the mitotic score at each amino acid position. The resulting difference score is close to zero when a position in the mitotic and flagellate sequences is equally conserved/diverged relative to the set of references sequences; it is positive when the mitotic sequence is less divergent, and negative when the mitotic sequence is more divergent. To identify the positions where the divergence of mitotic sequences was greater than flagellate sequences, the conservation score at each position was divided by the standard deviation of scores over all positions. We focused our subsequent analysis on especially divergent positions, which we defined as those where the relative divergence was greater than two standard deviations away from the mean (**Fig. 2A**).

We used PyMol (citation: The PyMOL Molecular Graphics System, Version 2.4.1 Schrödinger, LLC) and a cryo-EM structure of αβ-tubulin in a microtubule (PDB code 6O2R (Eshun-Wilson *et al.*, 2019)) to assess if the especially divergent positions in mitotic tubulins were enriched near microtubule polymerization interfaces (**Fig. 2B-C**, **Fig. S3**). To obtain the overall fraction of especially divergent positions per chain, the number of especially divergent positions in α- and β-tubulin was divided by the total number of amino acids. To calculate the proportion of divergent positions near lateral or longitudinal interfaces, we used distance based selections to identify the amino acids within a cutoff distance of a lateral or longitudinal lattice neighbor, and calculated the ratio of divergent to total positions within this subset.

### Cell and bacterial culture

*Naegleria* amoebae (strain NEG, ATCC strain 30223) and their food source *Aerobacter aerogenes* (a gift from the laboratory of Chandler Fulton, Brandeis University) were routinely cultured following previously established protocols (Heuser and Razavi, 1970). Briefly, *A. aerogenes* were regularly streaked from a frozen glycerol stock, and single colonies were grown stationary at room temperature in penassay broth (Difco antibiotic medium 3). Liquid cultures were used to grow lawns of *A. aerogenes* overnight on NM plates (2 g/L Difco Bacto peptone, 2 g/L glucose, 1.5 g/L K2HPO4, 1 g/L KH2PO4, 20 g/L agar). Lawns were inoculated with a loopful of NEG amoebae or cysts to create an edge plate (from a previous edge or cyst plate). Plates were sealed with parafilm, inverted, and incubated for 1-3 days at 28 °C. For starvation-induced differentiation (**Fig 1B**), cells were shocked with ice cold 2 mM Tris, and transferred to a shaking flask at 28 °C for 1 h.

### Mitotic synchronies

To obtain a population of synchronized cells, we modified a previously published method (Fulton and Guerrini, 1969) to cause a heat-induced mitotic arrest. Briefly, the day before the synchrony, a lawn of *A. aerogenes* was collected in 10 ml of TrisMg (2 mM Tris + 10 mM MgSO4), pelleted, resuspended in 20 ml TrisMg. 10 ml of the bacterial solution were transferred into a 125 ml flask. 2-8×10^5 amoebae were added to the flask and covered with foil, and the culture was incubated in a shaking water bath overnight (125 RPM, 30 °C). The morning of the synchrony, two additional lawns of *A. aerogenes* were collected, pelleted, and resuspended in 40 ml TrisMg. This solution was added to the flask with *Naegleria*, and allowed to shake for 3 minutes to thoroughly mix. This mixture was divided into 2 new (uncovered) flasks, one “control” and one “experimental,” and cell counts were taken with a hemocytometer. Cells were counted approximately every 20 min, and once the cells had doubled from their starting concentration, a sample was taken for quantitative real time PCR (qPCR) analysis (see next section), and the experimental flask was moved to a 38.5 +/− 0.5 °C water bath. Cells were counted from each flask, and when the control flask had doubled again, another sample was taken from each flask for qPCR, and then the experimental flask was shifted back to 30 °C. Samples were taken from the experimental flask after shifting back to 30 °C to fix and stain cells for mitotic spindles.

### Analysis of tubulin gene expression

Samples were collected from each flask prior to the temperature shift (pre-shift, control and experimental flasks), and again after incubation at 38 °C (or 30 °C for the control flask) but before shifting back to 30 °C. For each sample, 5 ml of cells were spun down at 1500 RCF at 4 °C for 5 min and the supernatant was discarded. The cell pellet was suspended in 1 ml TRIzol, vortexed, and promptly stored at −80 °C until RNA extractions. Cells were lysed using FastPrep homogenizer with bead beating in TRIzol. Lysate was cleaned up using a Zymo kit with on column DNase treatment, and RNA was eluted in 30 μl of kit-provided water. cDNA libraries were then generated using a Thermo Fisher/Invitrogen SuperScript™ IV First-Strand Synthesis System (Catalog #18091200). cDNA, PowerSybr Green (Thermo Fisher: 4368706), and primers were mixed in triplicate in a MicroAmp™ Fast Optical 96-Well Reaction Plate with Barcode (Catalog #4346906) and sealed with an optical adhesive cover (Catalog #4360954). Primer sequences were as follows: GAPDH (JGI ID: 53883): forward TGGCTCCAATTGCTGCTGTTT, reverse CCTTAGCAGCACCAGTTGAAGA; G protein (77952): forward ACGGTTGGGTCACTTGTTTGTCC, reverse GAGCGTGACCAGTGAGGGATC; mitotic α-tubulin (58607): forward GGTCCTTGATGTGTGCCGAAC, reverse TTAGCAGCATCTTCACGACCAGT; mitotic α- tubulin (55745) forward CACACACAAAATGAGAGAAGTCGTC, reverse TTCCATGTTCAGCACAGAATAATTC; mitotic β-tubulin (55748): forward AACCAACACTGCTTCTCCACTCG, reverse TCTGGACGGAATAATTGACCTTGG; mitotic β-tubulin (55900): forward GGTTGCTGGTGTCATGTCTGGTG, reverse GCAGCCAAAGGAGCAGAACCAA. Samples were run on a StepOne Real-Time PCR machine and analyzed using StepOne software v2.3.

The fold change in mRNA abundance was determined from C_T_ values using the 2^−ΔΔCt^ method (Livak and Schmittgen, 2001). Using this method, the flask that remained at 30 °C was a time-matched control for the experimental flask at the time point before the temperature shift, and the time point after the shift to 38 °C. A *Naegleria* G protein was used as the housekeeping gene to normalize the data, and a second housekeeping gene (GAPDH) was used to verify the results.

The microarray data in **Fig. 1D** was originally acquired in (Fritz-Laylin and Cande, 2010). Each experimental replicate had been completed with 2 technical replicates, so the technical replicates were first averaged. Then, the mRNA abundance at the 0 min time point (before differentiation) and at the 80 min time point (after differentiation to flagellates) were compared for each biological replicate to calculate the fold change in mRNA abundance for mitotic and flagellate tubulins.

### Immunofluorescence

Immunofluorescence staining of amoebae and flagellates in Fig 1B was performed using an actin cytoskeleton fixation protocol modified from (Velle and Fritz-Laylin, 2020). Cells were taken from an edge plate or from a sample of differentiated cells (see above), spun down at 1500 RCF for 90 sec, and cell pellets were resuspended in 1.5 ml 2 mM Tris. Cells were fixed in an equal volume of 2x fixative (50 mM sodium phosphate buffer, 125 mM sucrose, and 3.6% paraformaldehyde) for 15 minutes, then transferred to a 96 well glass-bottom plate coated with 0.1% poly(ethyleneimine) and allowed to settle for 15 min. Cells were rinsed twice in PEM (100 mM PIPES, 1 mM EGTA, 0.1 mM MgSO4; pH ~7.4) and permeabilized for 10 min in PEM + 0.1% NP-40 Alternative (Millipore, 492016) + 6.6 nM Alexa Fluor™ 488 Phalloidin (and 0.2x Tubulin Tracker Deep Red (Life Technologies, T34077, prepared according to manufacturer instructions) columns 1, 2 and 4 only). Cells were rinsed twice in PEM, then blocked in PEMBALG (PEM + 1% BSA, 0.1% sodium azide, 100 mM lysine, and 0.5% cold fish water gelatin; pH 7.4) at room temperature for 1 h. Cells were then incubated in primary antibody (anti-α-tubulin mouse monoclonal antibody (clone DM1A), Sigma, T6199) diluted to ~10 μg/ml in PEMBALG for 1 h. Cells were washed 3 times in PEMBALG, then incubated at room temperature for 1 h in Alexa Fluor™ 555 conjugated goat anti-mouse secondary antibody (Life Technologies, A21424) diluted to 2 μg/ml in PEMBALG, with 1x Tubulin Tracker Deep Red, ~66 nM Alexa Fluor™ 488 Phalloidin, and 1 μg/ml DAPI. Cells were then rinsed 4 times in PEM, and imaged the same day.

Immunofluorescence staining in the remaining figures was optimized for microtubules and performed using amoeba from a fresh edge plate that had grown about half-way across the dish (or from a mitotic synchrony, detailed above). Cells were removed from the plate and added to approximately 3 mls of water in a conical tube, spun down in a clinical centrifuge at setting 7 for ~40 seconds and the supernatant removed leaving ~500 μl of water above the cell pellet. To this mixture an equal volume of freshly prepared 2X fixative solution consisting of 2 mM Tris pH 7.2; 125 mM sucrose; 10 mM NaCl, 2% paraformaldehyde was added and mixed gently. Cells were fixed for 10 min at room temperature. Cells were then placed on freshly coated coverslips and allowed to adhere for approximately 20-30 minutes. Coverslips were plasma cleaned and then coated with 0.1% poly(ethyleneimine). After cells were adhered to the coverslips, they were rinsed 3 times with 1 ml of PEM (100 mM PIPES, pH 6.9; 1 mM EGTA; 0.1 mM MgSO4) and then permeabilized with 0.1% NP-40 for 10 minutes. Cells were blocked in PEM-BALG (PEM buffer supplemented with 1% BSA, 100 mM lysine, and 0.5% cold fish water gelatin) for one hour or overnight and then incubated with primary antibody for 1 hour at 37 °C or at room temperature overnight. Coverslips were rinsed in PBS containing 0.1% Tween and 0.02% sodium azide and incubated with Dylight-488 labeled anti-mouse secondary antibodies (Invitrogen) according to the manufacturers’ recommended protocol. Finally, coverslips were washed in PEM supplemented with 0.01% Triton-X-100 for 5 minutes before mounting on clean slides using DAPI Fluoromount G (Southern Biotech) or Prolong Gold.

### Confocal imaging

Cells were imaged on a Nikon Ti-E microscope with a CSU-X1 Yokogawa spinning-disk confocal scan head (PerkinElmer, Wellesley, MA), an Andor iXon+electron-multiplying charge-coupled device camera (Andor), using a 100X/1.4 NA objective lens. Z-step size was set at 0.2 μm.

Laser powers and exposures were chosen to ensure that the fluorescent signal would not be saturated and were adjusted depending on the fluorescent signal. For imaging microtubules with a Dylight 488 labeled secondary antibody, images were acquired using a 488 nm laser at 10.2% power; for imaging DNA, the 405 nm laser was used at 40.2% power.

The images in Fig. 1B were taken on a Nikon Ti2 microscope equipped with a Plan Apo λ 100x oil objective (1.45 NA), a Crest spinning disk (50 μm), a Prime 95B CMOS camera, and a Spectra III/Celesta light source (at 50-60% power with excitation wavelengths of 477, 546, and 638 nm). The microscope was controlled through NIS Elements software, and images were acquired as multi-channel z stacks with a step size of 200 nm and exposures of 200 ms (to image fluorescent phalloidin and tubulin antibody staining) or 500 ms (to image tubulin tracker staining).

### Digital deconvolution and 3D reconstructions

Z stacks captured using a spinning disk confocal microscope were digitally deconvolved using Autoquant X3 software. The default 3D deconvolution settings for spinning disk confocal data were used with “expert recommended settings,” and 40 iterations. The deconvolved images were then processed in Fiji (Schindelin *et al.*, 2012) to set the scaling, and to remove the mitochondria prior to 3D rendering, as the intensely-stained mitochondria made it difficult to observe the DNA in the nucleus. The resulting deconvolved image stacks were used to generate 3D surface renderings in UCSF ChimeraX software (Pettersen *et al.*, 2021).

### Analysis of spindle morphology

Spindle length and width measurements were assessed using the raw confocal (not deconvolved) datasets, and were only measured for spindles lying parallel to the plane of the coverslip. Length was measured by drawing a line in Fiji using the straight line tool, and measuring from the end of one pole to the opposite pole. For spindles in prophase where the poles are unclear, the longest axis was measured. In cases where the spindle bent during telophase (e.g. **Fig. 3A**, Anaphase/Telophase), the segmented line tool was used to follow the length of the spindle more accurately. Spindle width was measured using only the straight line tool, and was assessed at the approximate midpoint of the spindle between the two poles. These length and width values were separated by spindle stage, and were plotted using GraphPad Prism 8 software.

The number of bundles and the distance between bundles were calculated from confocal Z-stacks of metaphase spindles lying perpendicular to the coverslip. Bundle number was assessed in each plane going through the bundle for 8 representative spindles (**Fig. 4B**), and the maximum number of bundles present at the midplane was calculated for additional metaphase spindles. To determine the average distance between bundles, a frame that represented the spindle midplane was used, and the center of each bundle was selected using the multi-point tool in Fiji. The coordinates of each bundle center were used to determine the distance from each bundle to its two nearest neighboring bundles.

Line scan analysis (**Fig. 4D**, **Fig. S6**) was completed using confocal images of spindles that were oriented parallel to the coverslip. Image stacks were first transformed into sum intensity projections in Fiji. Then, the line width was matched to the width of the spindle, and a line (or segmented line in the case of bent anaphase/telophase spindles) was drawn to include the entire spindle length, with a short length of background at each end. The “plot profile” tool in Fiji was then used to extract the average pixel intensity along the line for tubulin and DNA staining. These values were normalized to the average intensity of an area of the cell adjacent to the spindle, which was set to 1. The spindle lengths were also normalized such that “0” represents the midpoint of the spindle. To determine the relative quantity of DNA and tubulin in these spindles (**Fig. 4E**), the area under the linescan-generated curves was calculated using GraphPad Prism 8 software, using a baseline level of 1.

### Analysis of spindle twist

To characterize the shape of microtubule bundles, we manually tracked individual bundles of vertically oriented spindles, and horizontally oriented spindles whose image stacks were first transformed into vertical (end-on) orientation, using Multipoint tool in Fiji. As microtubule bundles appear as spots in a spindle cross-section, each point was placed at the center of the signal and its x,y,z coordinates were saved. Moving up and down through the z-stack helped to determine this point. Each bundle was tracked through all z-planes where it was visible. Positions of the spindle poles were also determined, as the spots in the center of the end points of all bundles in the plane beyond the bundle ends. Coordinates of bundles and poles were transformed so that both poles are on the z-axis.

To describe the shape of a microtubule bundle, we fit a plane to the points representing the bundle. Subsequently, we fit a circle that lies in this plane to the same points. These fits were used to calculate the curvature and twist of the bundle as follows: (i) The curvature is calculated as one over the radius, and (ii) the twist is calculated as the angle between the plane and the z-axis divided by the mean distance of these points from the z-axis. Bundle length was calculated as the length of the projection of the bundle trace onto the pole-to-pole axis. For detailed descriptions of this method, see (Ivec *et al.*, 2021).

### Transmission Electron Microscopy

Cells were fixed overnight at 4 °C in 2.5% glutaraldehyde + 100 mM sodium cacodylate, then rinsed and stored in 100 mM sodium cacodylate overnight. Samples were then rinsed in 100 mM sodium cacodylate buffer, pH 7.4, three times for 10 minutes per wash. Cells were post fixed in 1% aqueous osmium tetroxide (Electron Microscopy Sciences) in 100 mM sodium cacodylate buffer overnight at 4 °C. Cells were then rinsed twice in water for 10 min per wash, before en bloc staining with 1% uranyl acetate (Electron Microscopy Sciences) in water for 1 hour at room temperature. Cells were rinsed 3 times in water, for 10 min per wash. Cells were then subjected to a graded ethanol dehydration series as follows with 15 min washes at each of the following ethanol concentrations: 50%, 70%, 80%, 90%, 95%, followed by two ten minute washes in 100% ethanol. Cells were quickly rinsed in propylene oxide, then infiltrated with 50% resin (Araldite 502/Embed-12, Electron Microscopy Sciences) and propylene oxide overnight. Cells were then incubated for 6-12 hours in each of the following resin concentrations: 70%, 85%, 95%, and 100% followed by embedding in 100% resin at 60 °C for 4 days. ~70 nm thin sections were cut using an RMC PowerTime XL Ultramicrotome with a Diatome diamond knife, and were transferred to copper grids. Sections were post stained with 1% uranyl acetate for 6 min, and lead citrate for 2 min. Images were taken using a JEOL JEM-200CX transmission electron microscope.

## ACKNOWLEDGEMENTS

We thank Nenad Pavin for discussions about the analysis of spindle twist, Shane Hussey for assistance with sequence analysis, Joshua Rafferty and Shadi Mahjoum for technical assistance, Laura Katz (Smith College) for assistance with the PhyloToL pipeline, and Sandra Balduaf (Uppsala University) for *Acrasis* tubulin sequences. We thank Chandler Fulton (Brandeis University) for strains and advice. We thank Alfredo Guzman for designing primers, Ravi Ranjan (University of Massachusetts, Genomics Resource Laboratory) for RNA extractions, and Madelaine Bartlett (UMass) and Courtney Babbitt (UMass) for qPCR equipment. We thank Andrew Kennard (UMass) and Tom Maresca (UMass) for comments on the manuscript. Light microscopy data was gathered in the Light Microscopy Facility and Nikon Center of Excellence at the Institute for Applied Life Sciences, UMass Amherst. We thank Kasia Hammar (Marine Biological Laboratory) for assistance with Transmission Electron Microscopy. 3D reconstructions were generated in UCSF ChimeraX, developed by the Resource for Biocomputing, Visualization, and Informatics at the University of California, San Francisco, with support from National Institutes of Health R01-GM129325 and the Office of Cyber Infrastructure and Computational Biology, National Institute of Allergy and Infectious Diseases. This work was supported by the National Institute Of Allergy And Infectious Diseases of the National Institutes of Health under Award Number F32AI150057 to K.B.V. and Award Number 1R21AI139363 to L.K.F.-L., and a Smith Family Foundation Award for Excellence in Biomedical Science to L.K.F.-L. The work of doctoral students M.T. and A.I. has been supported by the “Young researchers’ career development project – training of doctoral students” of the Croatian Science Foundation. M.T., A.I. and I.M.T. acknowledge the support of the European Research Council (ERC Consolidator Grant, GA Number 647077) and the Croatian Science Foundation (HRZZ project IP2019-04-5967). Work in L.M.R.’s lab was supported by the NSF (MCB-1615938 and MCB-2017687) and the Robert A Welch Foundation (I-1908).

