## Supplemental Figures for "*Naegleria’s* mitotic spindles are built from unique tubulins and highlight core spindle features"

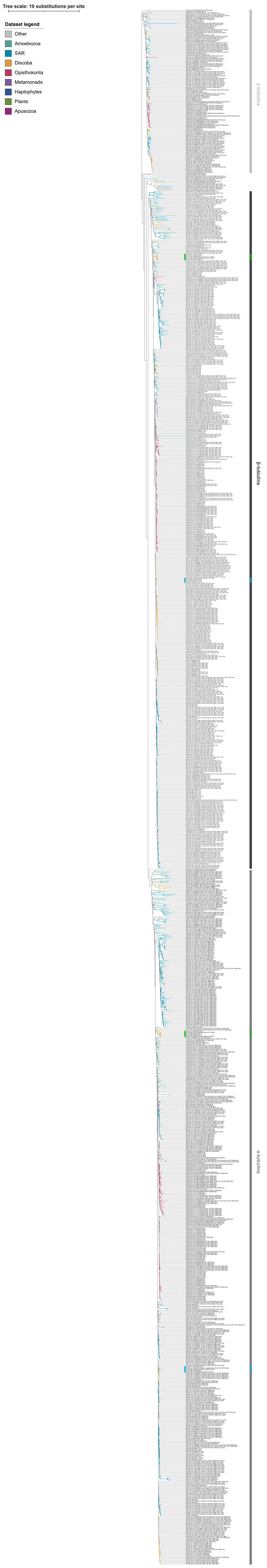


**Figure S1. Tubulin phylogeny.** A phylogram of 1,191 tubulins from 200 different species across the tree of life. Branches are colored according to the overarching group of the species represented by each sequence: teal, Amoebozoa; light blue, SAR; orange, Discoba; red, Metazoa; purple, Metamonads; dark blue, Haptophytes; green, Plants; maroon, Apusozoa; grey, Other. Blue vertical lines indicate flagellate tubulins, and green indicates mitotic tubulins. Symbols at the tips of branches indicate tubulins from *Acrasis* (open circles) or *Naegleria* (closed circles).

**A**

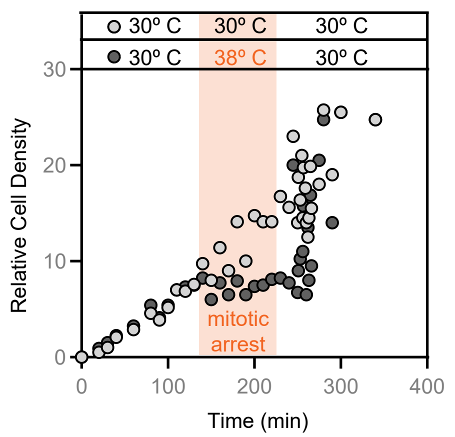


**B**

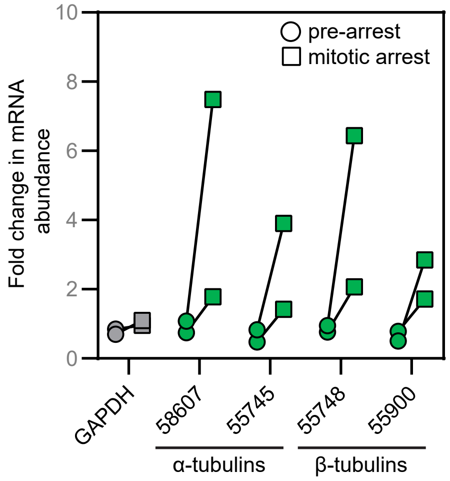


**C**

**α-tubulins**

| | | | | | flagellate α-tubulins (heteroloboseans) | | | | mitotic α-tubulins (heteroloboseans) | | | | |
|---|---|---|---|---|---|---|---|---|---|---|---|---|---|
| | *H. sapiens* (ENSP-00000336799) | *C. reinhardtii* (186023) | *S. cerevisiae* (288cYML085C) | *S. cerevisiae* (288cYML124C) | *N. gruberi* (53284) | *N. gruberi* (56236) | *A. kona* (12591) | *N. fowleri* (3292) | *N. gruberi* (58607) | *N. gruberi* (55745) | *A. kona* (12838) | *N. fowleri* (5134) | *N. fowleri* (7486) |
| *N. fowleri* (7486) | 57 | 60 | 55 | 55 | 60 | 61 | 60 | 61 | 71 | 88 | 67 | 73 | 100 |
| *N. fowleri* (5134) | 55 | 58 | 53 | 53 | 57 | 57 | 57 | 57 | 83 | 71 | 64 | 100 | |
| *A. kona* (12838) | 63 | 65 | 60 | 60 | 63 | 65 | 65 | 64 | 65 | 66 | 100 | | |
| *N. gruberi* (55745) | 58 | 62 | 56 | 55 | 61 | 62 | 63 | 62 | 73 | 100 | | | |
| *N. gruberi* (58607) | 57 | 59 | 54 | 54 | 58 | 59 | 59 | 59 | 100 | | | | |
| *N. fowleri* (3292) | 85 | 92 | 68 | 67 | 88 | 98 | 97 | 100 | | | | | |
| *A. kona* (12591) | 84 | 92 | 69 | 67 | 87 | 97 | 100 | | | | | | |
| *N. gruberi* (56236) | 85 | 92 | 68 | 67 | 88 | 100 | | | | | | | |
| *N. gruberi* (53284) | 79 | 86 | 68 | 67 | 100 | | | | | | | | |
| *S. cerevisiae* (288cYML124C) | 72 | 69 | 88 | 100 | | | | | | | | | |
| *S. cerevisiae* (288cYML085C) | 74 | 69 | 100 | | | | | | | | | | |
| *C. reinhardtii* (186023) | 86 | 100 | | | | | | | | | | | |
| *H. sapiens* (ENSP00000336799) | 100 | | | | | | | | | | | | |

**β-tubulins**

| | | | | flagellate β-tubulins (heteroloboseans) | | | mitotic β-tubulins (heteroloboseans) | | | | |
|---|---|---|---|---|---|---|---|---|---|---|---|
| | *H. sapiens* (ENSP-00000217133) | *C. reinhardtii* (129876) | *S. cerevisiae* (288cYFL037W) | *N. gruberi* (83350) | *A. kona* (08655) | *N. fowleri* (5358) | *N. gruberi* (55748) | *N. gruberi* (55900) | *A. kona* (11976) | *N. fowleri* (5966) | *N. fowleri* (3784) |
| *N. fowleri* (3784) | 57 | 63 | 62 | 63 | 63 | 63 | 70 | 88 | 67 | 71 | 100 |
| *N. fowleri* (5966) | 56 | 59 | 57 | 60 | 60 | 60 | 86 | 71 | 62 | 100 | |
| *A. kona* (11976) | 60 | 66 | 64 | 67 | 67 | 67 | 62 | 65 | 100 | | |
| *N. gruberi* (55900) | 58 | 63 | 62 | 63 | 64 | 64 | 71 | 100 | | | |
| *N. gruberi* (55748) | 57 | 60 | 58 | 61 | 61 | 60 | 100 | | | | |
| *N. fowleri* (5358) | 74 | 90 | 72 | 97 | 96 | 100 | | | | | |
| *A. kona* (08655) | 73 | 91 | 72 | 96 | 100 | | | | | | |
| *N. gruberi* (83350) | 75 | 91 | 72 | 100 | | | | | | | |
| *S. cerevisiae* (288cYFL037W) | 65 | 70 | 100 | | | | | | | | |
| *C. reinhardtii* (129876) | 73 | 100 | | | | | | | | | |
| *H. sapiens* (ENSP00000217133) | 100 | | | | | | | | | | |

% Identity

0 — 100

**Figure S2. The *Naegleria* tubulins expressed during mitosis are divergent.** (**A**) To enhance the percent of mitotic cells, a population of cells grown at 30 °C was shifted to 38 °C (dark gray points, orange panel) to induce a mitotic arrest. After shifting cells back to 30 °C, cells divide synchronously, quickly catching up to the density of cells in a control flask left at 30 °C (light gray points). Each point represents the relative cell density of a flask at a given time point (for 9-18 timepoints per flask), for 5 independent synchrony experiments. (**B**) Samples of cells from a mitotic synchrony experiment were subjected to qPCR analysis to determine mRNA levels from housekeeping genes (GAPDH, gray, and G protein, used for normalization), and mitotic α and β tubulins (green). The fold change in mRNA abundance before (circles) or after (squares) the 38 °C mitotic arrest was calculated relative to a control flask kept at 30 °C. Each point represents one biological replicate consisting of 3 technical replicates. (**C**) The percent identities of α (top) and β (bottom) tubulin sequences were calculated for different tubulins in multiple species. Purple indicates lower % ID, while gray indicates a higher % ID.

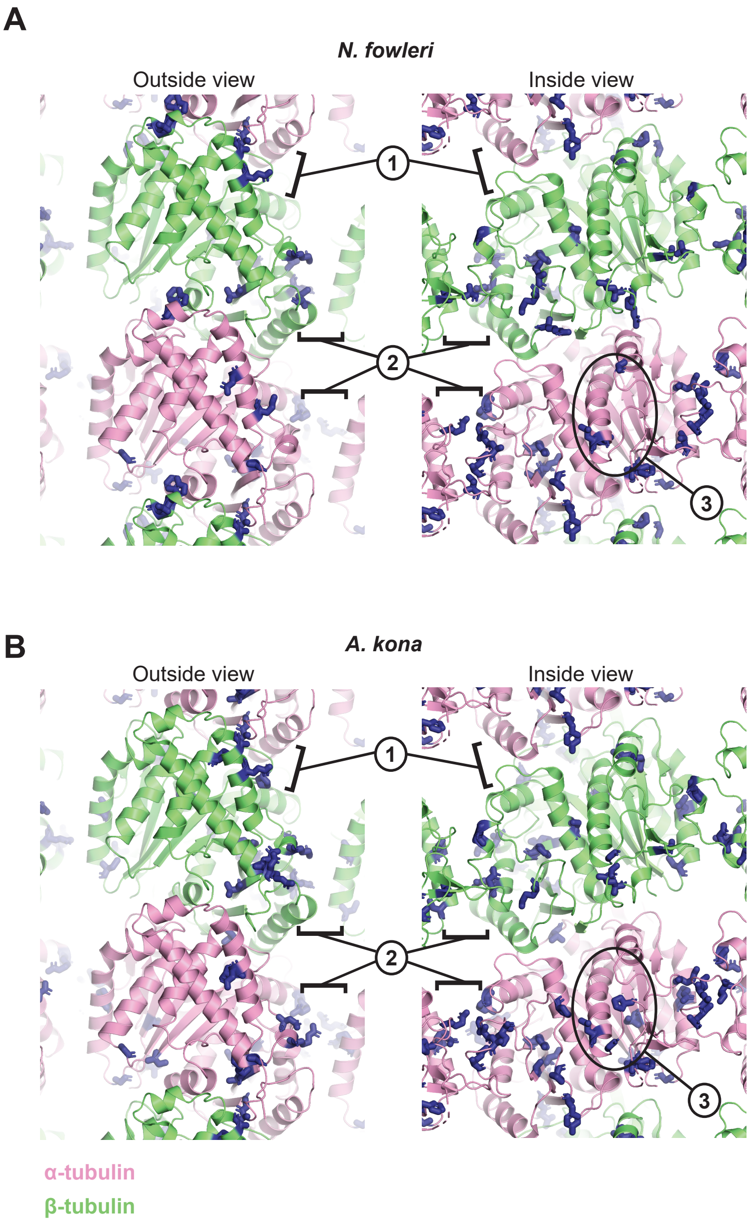


**Figure S3. Structural context of the sites with increased divergence in the mitotic tubulins.** (**A**) Side-chain positions for the *N. gruberi* amino acids identified in Fig. 2A are represented as sticks (blue) on a model of αβ-tubulin in the microtubule lattice (α-tubulin: pink, β-tubulin: lime). 'Outside' and 'Inside' views of the lattice are shown, and longitudinal (labeled 1) and lateral (labeled 2) microtubule lattice contacts are indicated, as is the luminal (internal) surface of α-tubulin (labeled 3). (**B**) As in A, but for the side-chain positions for the *A. kona* amino acids identified in Fig. 2A.

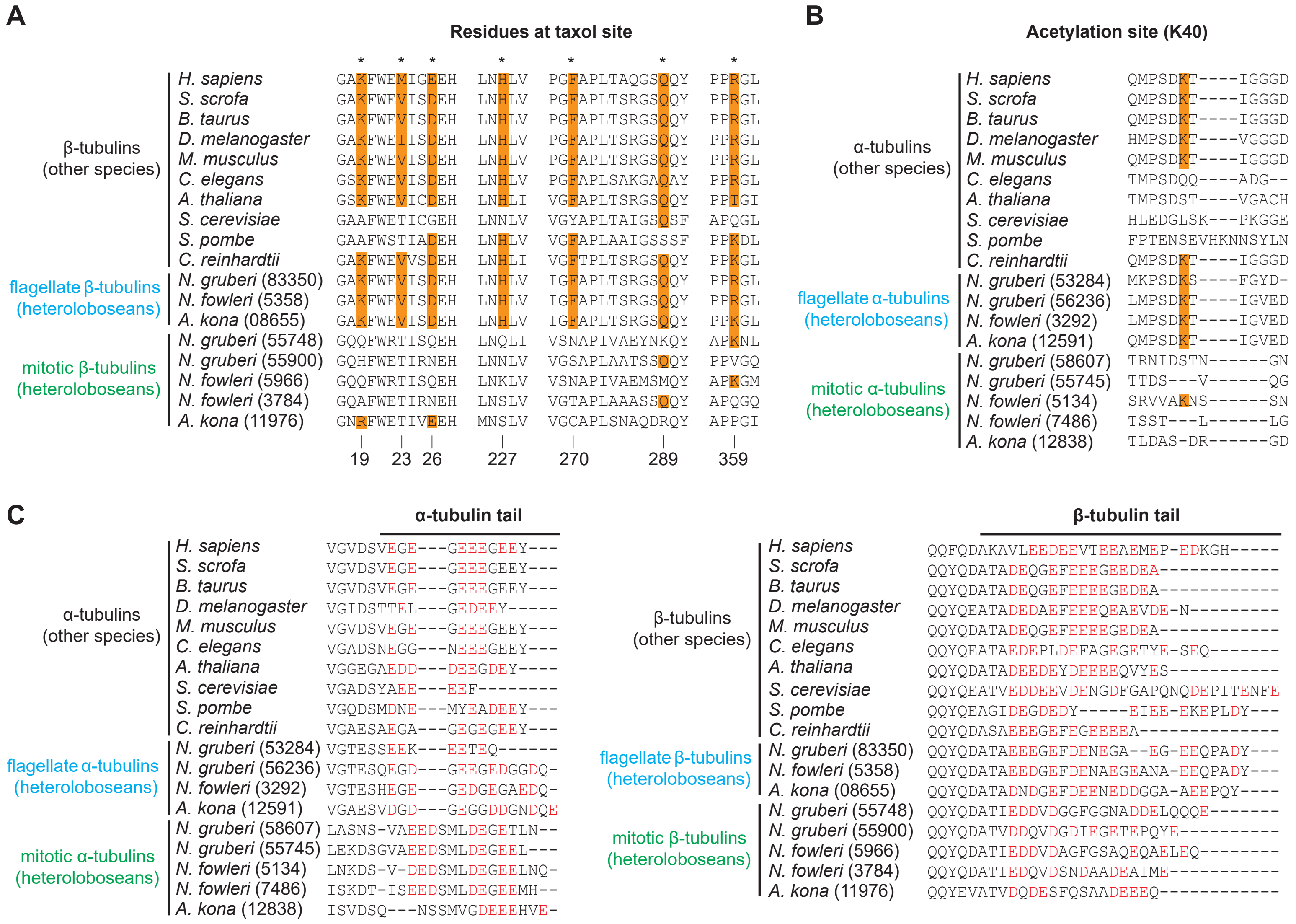

**Figure S4. Key differences between mitotic and flagellate sequences.** (**A**) Taxol binding: flagellate β-tubulins have generally conserved residues implicated in taxol binding (highlighted orange), but the mitotic β-tubulins have not. (**B**) α-tubulin acetylation: flagellate α-tubulins have generally conserved a K40-equivalent residue (highlighted in orange); this position is subject to acetylation in more commonly studied organisms. Mitotic α-tubulins have diverged in this region. (**C**) Tubulin 'tails': the mitotic α-tubulins are notably lacking the C-terminal tyrosine (Y) that is subject to detyrosination/retyrosination in more commonly studied organisms. Besides that, there are few obvious differences in the length or overall charge (negatively charged amino acids are colored red) between mitotic and flagellate α- or β-tubulins.

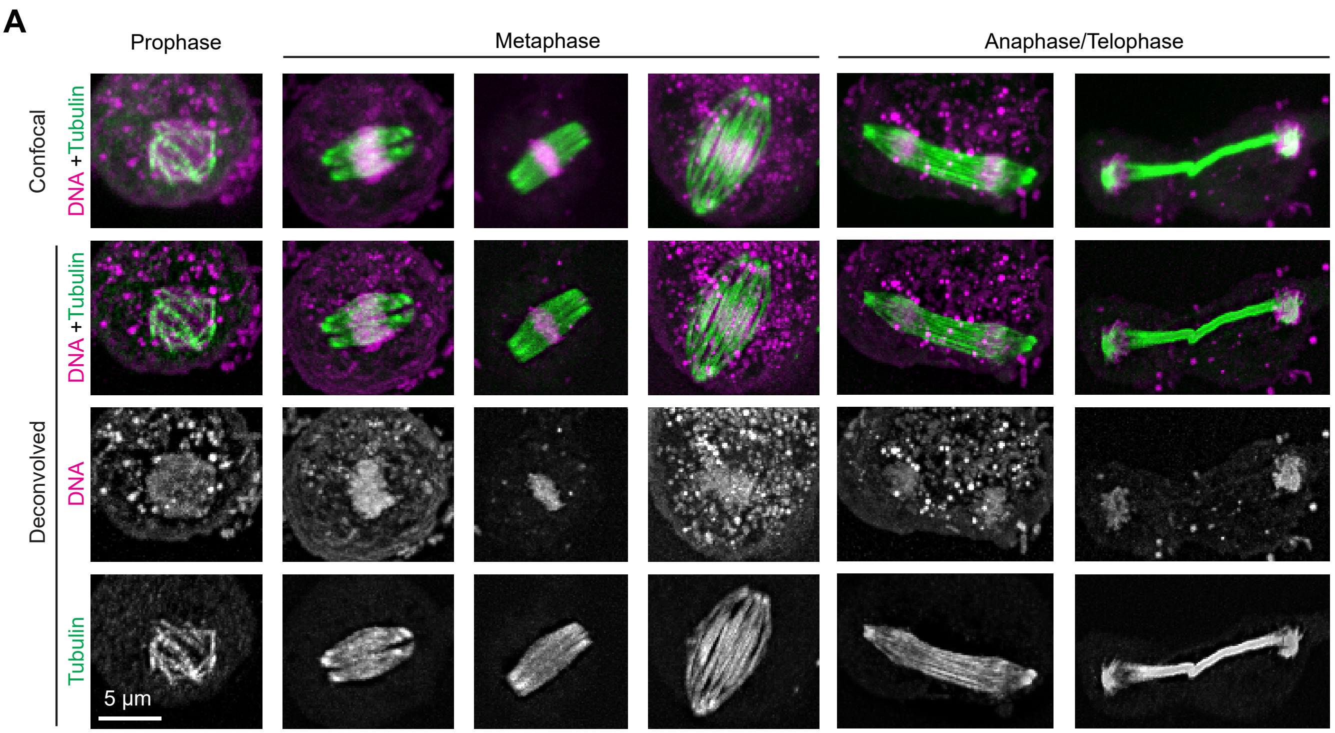


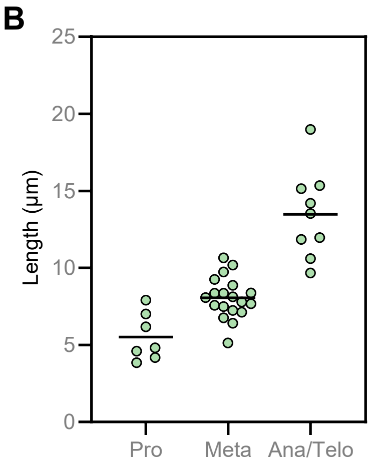


C

| **Prophase** | *Non-synched* | *Synched* | *All* |
|---|---|---|---|
| # | 6 | 7 | 13 |
| Length (µm) | 5.3±1.1 | 5.5±1.5 | 5.4±1.3 |
| Width (µm) | 4.6±0.5 | 4.2±1.1 | 4.4±0.8 |
| **Metaphase** | | | |
| # | 10 | 19 | 29 |
| Length (µm) | 6.3±0.6 | 8.1±1.3 | 7.4±1.4 |
| Width (µm) | 3.7±0.3 | 3.7±1.1 | 3.7±0.9 |
| **Ana/Telophase** | | | |
| # | 4 | 9 | 13 |
| Length (µm) | 16.5±3.7 | 13.5±2.8 | 14.4±3.3 |
| Width (µm) | 0.9±0.3 | 2.5±1.3 | 2.0±1.3 |

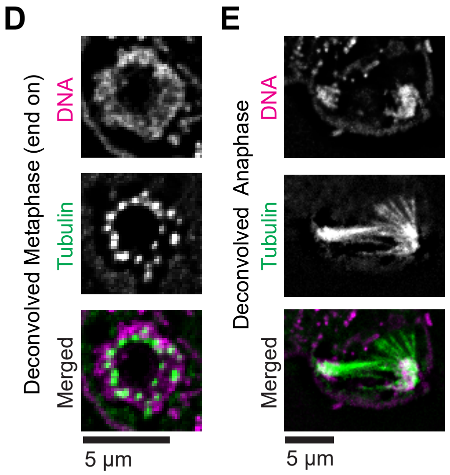


**Figure S5. Mitotically synchronized amoebae have indistinguishable spindle architecture from unsynchronized cells.** (**A**) Amoebae were synchronized, then fixed and stained with antibodies (anti-alpha tubulin clone DM1A, green) to detect microtubules, and DAPI to label DNA (magenta). Cells were imaged using confocal microscopy (top row), and the images were deconvolved using Autoquant software (bottom rows). All images are maximum intensity projections. Cells were classified as prophase, metaphase, or anaphase/telophase. (**B**) Cells were treated as in A, and confocal images were used to quantify the maximum spindle length. Each point represents one mitotic spindle, and lines indicate the averages. (**C**) Spindle lengths and widths were measured for spindles of each stage, and either grouped by methodology (synched vs non-synced) or pooled (all). The number of spindles measured is shown (#), and average lengths and widths are given with SD. (**D**) A spindle (from experiments like those in A) lying perpendicular to the coverslip was imaged using confocal microscopy, and deconvolved. The image represents a single z plane at the midpoint of the spindle. (**E**) A cell (from experiments like those in A) with an apparent spindle irregularity reveals each bundle may be composed of multiple microtubules. A single z plane is shown.

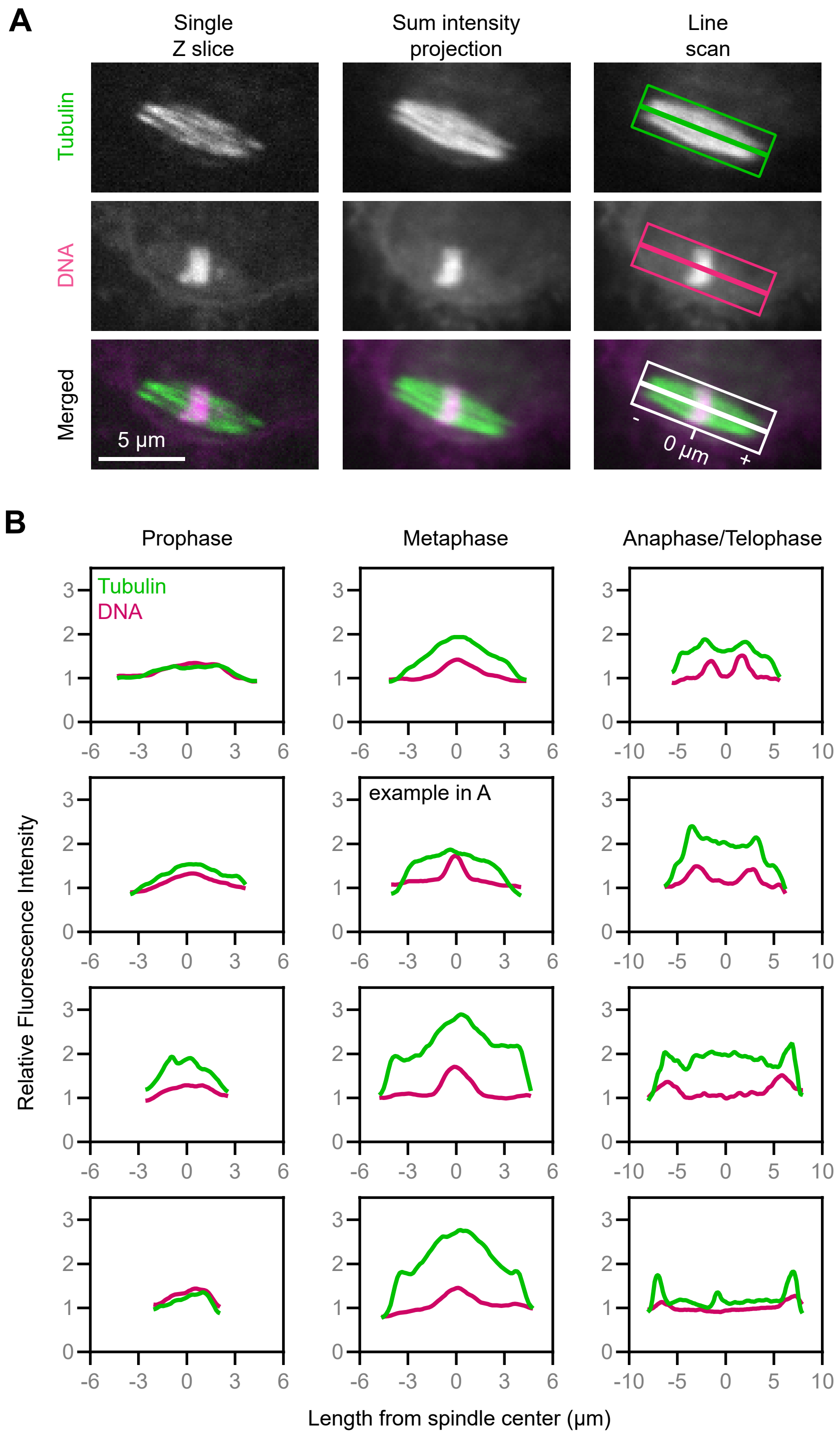


**Figure S6. Additional examples of line scan analysis.** (**A**) Populations of synchronized cells were fixed and stained for tubulin (green) and DNA (magenta), and cells lying in the plane of the coverslip were imaged using a spinning disk confocal microscope. Sum intensity projections (center panels) were generated in Fiji, and line scans (right panels) were drawn from pole to pole (thick line), with a line thickness adjusted to encompass the entire spindle width (thinner box). The center of the spindle was set to 0 μm. (**B**) Line scans were drawn on spindles in prophase (left), metaphase (center), and anaphase/telophase (right). The pixel intensities along the spindle length were normalized to the average intensity of an area in the cell adjacent to the spindle, which was set to 1.

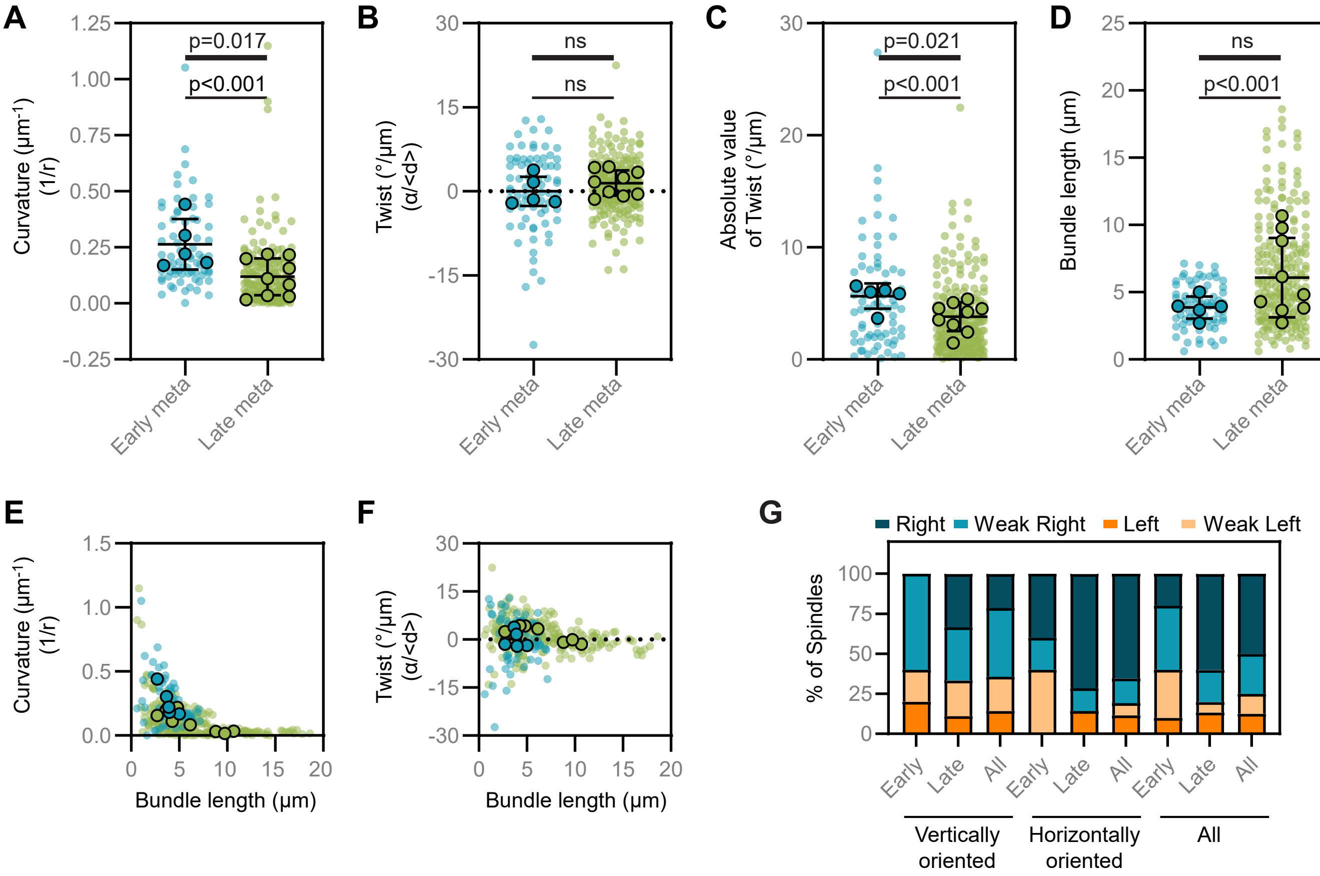


**H**

| Stage | Curvature (1/µm) | Twist (°/µm) | Bundle length (µm) | Spindle length (µm) | Number of bundles | Number of spindles |
|---|---|---|---|---|---|---|
| **Early metaphase** | 0.250 ± 0.021 | 0.059± 0.847 | 3.96 ± 0.18 | 7.19 ± 0.78 | 75 | 5 |
| **Late metaphase** | 0.111 ± 0.009 | 1.143 ± 0.313 | 6.43 ± 0.26 | 14.55 ± 2.56 | 226 | 9 |
| **All metaphase** | 0.146 ± 0.009 | 0.873 ± 0.316 | 5.81 ± 0.21 | 11.92 ± 1.90 | 301 | 14 |

**Figure S7. Quantification of the curvature and twist of microtubule bundles.** (**A**) Curvatures were calculated for individual bundles (smaller data points) and averaged for each spindle (larger data points). Lines indicate the mean and standard deviation calculated from spindle averages. Early metaphase bundles are significantly more curved when analyzed per bundle (indicated by thin line, Mann-Whitney test), or when spindle averages are compared (indicated by thick line, unpaired t test). (**B**) Bundle twists were calculated and compared, and are displayed as in A. The mean twist is different from 0 in late metaphase ($p = 0.0003$), but not in early metaphase ($p = 0.94$). To determine whether spindles are statistically different in early versus late metaphase, we compared the twist of both individual bundles and whole spindles and found no statistically supported difference (individual bundles: $p = 0.233$, unpaired t test, whole spindles: $p = 0.325$, unpaired t test). (**C**) The twist values in panel B were converted to absolute values. These values support more total twist in early metaphase than late for individual bundles (Mann-Whitney test) and whole spindles (unpaired t test). (**D**) Bundle lengths were measured and data are displayed for individual bundles and spindle averages as in A. Late metaphase spindles have longer bundles when all individual bundles are considered (Mann-Whitney test), but not when averaged by spindle ($p = 0.061$, unpaired t test). (**E**) The curvature of microtubule bundles is shown as a function of bundle length. Each small dot represents a single bundle within a spindle, while each larger dot represents the average for a spindle. Teal dots indicate bundles within early metaphase spindles, while green dots indicate late metaphase. (**F**) The twist of microtubule bundles is shown as a function of bundle length. Each small dot represents a single bundle within a spindle, while each larger dot represents the average for a spindle. Teal small dot represents a single bundle within a spindle, while each larger dot represents the average for a spindle. Teal dots indicate bundles within early metaphase spindles, while green dots indicate late metaphase. (**G**) The percentage of spindles with right, weak right, left, or weak left handedness are shown. Spindles are grouped according to the stage of mitosis (early or late metaphase) and their orientation (vertical or horizontal), as indicated. The twist was determined visually by moving through end-on z-stacks from the bottom plane towards the top plane, where bundle rotation clockwise and counterclockwise implies a left-handed and right-handed twist, respectively. For horizontally oriented spindles, z-stacks were first rotated to obtain the end-on view. (**H**) The mean values (+/- SEM) for the bundle characteristics quantified above are shown.
